# Cognitive Mode Detectable with Task-Based fMRI: Default Mode B (DMB)

**DOI:** 10.1101/2025.04.21.649745

**Authors:** Yi Qing Yvette Ni, Solana Redway, Aida Momeni, Ava Momeni, Lujia Jian, Lily Yip, Todd S. Woodward

## Abstract

In the context of task-based functional magnetic resonance imaging (fMRI), cognitive modes can be defined as task-general cognitive/sensory/motor processes which reliably elicit specific blood-oxygen-level-dependent (BOLD) signal pattern configurations. A number of cognitive modes are detectable with task-based fMRI, and here we focus on Default Mode B (DMB), a task-negative and late-trial peaking cognitive mode. The BOLD signal configurations associated with DMB are modulated by a range of tasks, and here we present eight. For each task, we report: (1) specific pattern-based (as opposed to coordinate-based) anatomical details essential for distinguishing DMB from other BOLD-based cognitive modes, and (2) task-induced BOLD signal changes associated with DMB over a range of task conditions. In order to facilitate recognition, we nick-named the anatomical patterns specific to DMB as follows: (1) In Flight, (2) Medial Temporal Dots, (3) Snowman Nose, (4) Angel Wings, and (5) Tripod. Evidence for DMB was derived from the timing and magnitude of task-induced BOLD signal changes induced by the following tasks: working memory, spatial capacity, semantic association, evidence integration, Raven’s matrices, autobiographical event simulation, meditation and social perception. It was observed that deactivations in DMB were sensitive to cognitive load during attention to specific features of the external environment, based on evidence from working memory, spatial capacity, semantic association, evidence integration, and Raven’s matrices. It was also observed that activations in DMB involved a cognitive process for engaging in mental projection into self-relevant social narratives, based on evidence from autobiographical event simulation, meditation, and social perception. Future research may explore DMB activation over a wider range of tasks in larger samples.

## 1. Introduction

In the context of task-based functional magnetic resonance imaging (fMRI), cognitive modes can be defined as task-general cognitive/sensory/motor processes which reliably elicit specific blood-oxygen-level-dependent (BOLD) signal pattern configurations (Evora & Woodward, 2025; Jian et al., 2024; Mascarenhas et al., 2024; Momeni et al., 2024; Percival et al., 2025; Redway et al., 2024; Redway et al., 2025; Wang et al., 2024; Zeng et al., 2024). How these cognitive-mode BOLD patterns overlap anatomically with resting state networks has been put forward in a number of publications (Fouladirad et al., 2022; Gill et al., 2022; Kusi et al., 2022; Momeni et al., 2025; Sanford et al., 2020a, 2020c; Sanford & Woodward, 2021a, 2021b; Wong et al., 2020; Zurrin et al., 2024). However, as opposed to resting-state fMRI, task-based fMRI allows observation of task-induced BOLD signal changes over a range of experimental conditions, and inspection of these changes is how the cognitive mode associated with each anatomical configuration can be identified and supported.

A number of cognitive modes are detectable with task-based fMRI (Chen & Woodward, 2024; Percival, Chen, & Woodward, 2024; Percival, Chen, Zahid, et al., 2024; Percival et al., 2020), with a mid-trial peaking cognitive mode being the focus here, referred to as Default Mode B (DMB). The goals of the current study were to: (1) determine the replicability of DMB and its associated anatomical patterns, and (2) provide evidence for the DMB cognitive mode by way of inspection of the task-induced BOLD signal changes over a range of fMRI tasks. This pattern-based methodology is an alternative to the traditional coordinate-based methodology (Lancaster et al., 2000), and provides an efficient way to communicate the configuration of hundreds or thousands of voxels with a single pattern. The DMB signature patterns show bilateral activity in the superior frontal gyri (BA 9), occipital cortex (BA 39), temporal middle gyri (BA 21), precuneous cortex (BA 23) and frontal medial cortex (BA 11). In order to facilitate recognition, we nick-named the anatomical patterns specific to DMB as follows: (1) In Flight, (2) Medial Temporal Dots, (3) Snowman Nose, (4) Angel Wings, and (5) Tripod, and how these link to the aforementioned anatomical regions are presented in detail below.

The process of identification of fMRI-specific cognitive modes is rooted in the “objective discovery” approach for determination of brain function, which calls on neuroscientists to embrace an objective field-specific vocabulary to replace/refine a vocabulary inherited from philosophy and psychology (Buzsaki, 2020). In the case of DMB, this could include avoiding the assumption that fMRI can detect the cognitive processes which the researchers had in mind when designing the task, including terms such as daydreaming, mind-wandering, and engaging in autobiographical memory. Although, these cognitive operations are clearly required for the tasks presented here, they need not be good definitions of the cognitive modes to which fMRI is sensitive during these tasks. Replacement of vocabulary inherited from philosophy and psychology with fMRI-specific cognitive modes provides a pathway to reverse-inference (Poldrack, 2006), with the end goal being a one-to-one mapping between a set of anatomical patterns on one hand, and cognitive modes on the other. Here we study reliable associations between anatomy and function in DMB through re-analysis of a series of task-based fMRI data collected in our laboratory and others’ from published and unpublished results.

## 2. Methods

The set of task-based cognitive modes and whole-brain anatomical patterns referenced here (Percival et al., 2020) were developed through a series of analyses using constrained principal component analysis for fMRI (fMRI-CPCA) (Metzak et al., 2011; Woodward et al., 2013). An overview of CPCA methodology has been published elsewhere (Hunter & Takane, 2002; Takane & Hunter, 2001; Takane & Shibayama, 1991) and its application to fMRI has been published in a series of works (Fouladirad et al., 2022; Gill et al., 2022; Kusi et al., 2022; Metzak et al., 2011; Sanford et al., 2020a, 2020c; Sanford & Woodward, 2021a, 2021b; Wong et al., 2020; Woodward et al., 2006; Woodward et al., 2013; Zurrin et al., 2024), and was summarized in an Organization for Human Brain Mapping (OHBM) Neurosalience podcast in May, 2024 (Woodward, 2024). Prototypical examples of 11 cognitive modes and whole-brain anatomical patterns were identified via study of specific activity patterns, and template images were created for each cognitive mode by anatomically averaging over these prototypical examples (Percival, Chen, Zahid, et al., 2024; Percival et al., 2020). Following this, a MATLAB-based algorithm was developed to classify newly derived brain images into these 11 task-based cognitive mode templates (Woodward et al., 2021). The algorithm assigns a *Z*-score to a to-be-classified brain image, reflecting how well activity matches the templates across the set of specific patterns.

### 2.1. Constrained Principal Component Analysis for fMRI (fMRI-CPCA) Overview

Typically, task-based fMRI studies use subtraction methodology, attempting to isolate cognitive-process-related brain activity by subtracting out the brain activity associated with cognitively simpler tasks. For example, the task control condition might exclude a cognitive process thought to be present in the task experimental condition. The subtraction process typically involves observing differences between beta weights derived from regressing the observed BOLD signal fluctuations onto a synthetic model of the hemodynamic response (HDR) pattern expected for the task timing built into the experimental design (Friston & Stephan, 2007). This is carried out at each voxel to derive a map of which voxels significantly differ with respect to their match to the assumed/synthetic HDR. However, this subtraction methodology for fMRI has limitations (Chinchani et al., 2025). First, subtraction is sensitive to how well task-induced BOLD changes match an assumed HDR but neglects direct observation of the task-induced BOLD changes elicited by the task conditions, when it is these observed task-induced BOLD changes are essential for recognition of cognitive modes. It’s plain to see that observed task-induced BOLD changes are unlikely to exactly match the assumed model shape, and as will be made clear below, they may differ substantially. Second, multiple cognitive processes (e.g., sustained attention, response, response re-evaluation) could all partially match a synthetic HDR pattern model, conflating them anatomically and interpretationally (Sanford et al., 2020c; Woodward et al., 2013). Last, each subtraction produces a single brain activation map, but this map can be a mixture of multiple cognitive modes which can only be separated using a dimensional approach to analysis.

An alternative to subtraction methodology for whole-brain task-based fMRI analysis with the goal of finding specific anatomical patterns is a dimensional analysis method that (1) does not require parcellating the brain into discrete regions, (2) integrates task trial timing information into the analysis in advance of extraction of dimensions, and (3) does not require matching to an assumed HDR shape to produce anatomical images. This can be achieved using a finite impulse response (FIR) model combined with a dimensional analysis constrained to task timing-predictable BOLD signal variance. The FIR model can be designed to produce an estimate of task-induced BOLD changes for each task condition, brain image, and participant separately. For fMRI studies, this leads to the separation of functional brain images onto dimensions, and interpretation of the cognitive modes associated with each image through inspection of task-induced BOLD changes (Metzak et al., 2011; Woodward et al., 2013).

In the current study, we use a FIR-based, dimensional analysis referred to as fMRI-CPCA (software code available at https://www.nitrc.org/projects/fmricpca) (Metzak et al., 2011; Woodward et al., 2013). Conceptually, fMRI-CPCA combines multivariate multiple regression and principal component analysis (PCA) into a unified framework. Multivariate multiple regression isolates variance in BOLD signal predictable from the independent variables, which are a set of FIR variables collected into a design matrix (Henson & Friston, 2007). Following this, PCA is performed on the matrix of BOLD signal regression-predicted scores (Pedhazur, 1997, p. 19, equation 2.9), derived from regressing the BOLD signal matrix onto the FIR model.

The fMRI-CPCA component loadings provide spatial information (BOLD-based anatomical patterns), which can be overlaid on a brain image. Computations based on the component scores provide temporal information. The component scores provide a value indexing the activity of each whole brain anatomical pattern for every full brain scan (repetition time; TR). When the component scores are regressed onto the FIR model matrix, predictor weights are obtained for every combination of participant, task condition and whole-brain anatomical pattern, due to the nature of the columns of the FIR model in the design matrix. It is these predictor weights that are plotted to produce task-induced BOLD signal changes. Thus, through fMRI-CPCA, we can: (1) identify multiple cognitive modes that are simultaneously involved in a cognitive task, (2) estimate the task-induced BOLD signal for each cognitive mode separately for each participant and condition, and (3) statistically test the effect of task conditions on estimated task-induced BOLD signal changes using standard repeated-measures analyses of variance (RM-ANOVAs). Structural and functional images were preprocessed using Statistical Parametric Mapping 8 (SPM8), or SPM12.

### 2.2. Anatomical Classification

#### 2.2.1. Template Image Creation

To date, whole-brain images have been created for each of the fMRI task-based cognitive modes (Percival et al., 2020) by averaging the images of several fMRI-CPCA components demonstrated to be excellent anatomical examples of a certain cognitive mode, in that they displayed distinctive and uniform anatomical and temporal activity patterns (Chen & Woodward, 2024; Percival, Chen, & Woodward, 2024; Percival, Chen, Zahid, et al., 2024).

#### 2.2.2. Component Classification

The template matching algorithm (Woodward et al., 2021) provides automatic classification information for any to-be-classified anatomical image to 11 task-based anatomical template images. Each of these 11 template images has its own unique set of 20-30 brain slices that highlight their replicable activity patterns (Chen & Woodward, 2024; Percival, Chen, & Woodward, 2024; Percival, Chen, Zahid, et al., 2024). For each classification, the algorithm isolates these specific slices from the template and the to-be-matched image. The values in the template image and the component loadings in the to-be-classified image are correlated using Pearson’s *r* and are then transformed to Fisher *Z* values. This is repeated for each template. This results in eleven Fisher *Z* values indicating how well the to-be-matched image “matches” each of the templates.

To retrieve the DMB examples reported here, 321 full-brain images from 74 fMRI-CPCA analyses with different combinations of 27 tasks were matched to the 11 templates. The top 8 matches for DMB were selected for presentation here (*Z* statistics in brackets): Working Memory with Thought Generation Task (WML46D04-TG; *Z* = 1.77); Spatial Capacity Task (SCAP; *Z* = 0.92), Semantic Association (SA; *Z* = 0.77), Picture Completion Bias Against Disconfirmatory Evidence (PCBADE; *Z* = 1.09); Raven’s Matrices (RSPM; *Z* = 0.80), Autobiographical Event Stimulation (AES; *Z* = 0.91), Mindfulness Meditation Task (MM; *Z* = 0.81), and Human Connectome Project: Social Task (HCP-SOC; *Z* = 0.91).

Two of the eight DMB anatomical images presented here were part of the 10 original anatomical images averaged to create the 2020 DMB image anatomical template (Percival et al., 2020). Specifically, the images from the WML46D04-TG task (Sanford et al., 2020c) and the AES task (Momeni et al., 2025) presented here were among the 10 images used to create the 2020 DMB anatomical template image. Specifically, scans of 27 participants completing the TG task were included twice in the template image.

In contrast, the SCAP (Sanford, 2019), SA (Woodward et al., 2015), PCBADE (Lavigne, Menon, Moritz, et al., 2020), RSPM (Zurrin et al., 2024), MM, and HCP-Soc images presented here provide entirely independent replications of the DMB pattern-based anatomy, as they were not in any way included in or overlapping with the original 2020 anatomical template image (Percival et al., 2020). All anatomical labels used in this paper are based on the Automated Anatomical Labeling (AAL) atlas included in MRIcron (Rorden & Brett, 2000) unless otherwise specified as Harvard Oxford atlas labels, and all X Y Z coordinates are in MNI anatomical space. WML46D04-TG, PCBADE, SA, RSPM and MM were pre-processed with SPM 8, and SCAP, AES and HCP-Soc with SPM 12.

### 2.3. Repeated Measures Analysis of Variance (RM-ANOVA) of Predictor Weights

fMRI-CPCA produces predictor weights for each component for each combination of post-stimulus time bin, task condition, and participant. These weights reflect the engagement of the whole-brain anatomical pattern at each trial time point, and provide an estimate of the task-evoked BOLD change, plotted over time, averaged over trials. This allowed BOLD change activity in each of the cognitive modes to be statistically examined by applying repeated measures analyses of variance (RM-ANOVAs) to the predictor weights. The Greenhouse-Geisser correction was applied when the assumption of sphericity was violated, and the uncorrected degrees of freedom were reported only when statistical significance was achieved using the Greenhouse-Geisser corrected degrees of freedom. Inspection of the task-evoked BOLD changes while considering the experimental designs provided the definition of the DMB cognitive mode.

Due to the complexity of interpreting interactions with within-subjects variables having many levels, the significant interactions were analyzed by comparing the BOLD changes at adjacent time points. That is to say, Time × Condition interactions were interpreted as sets of 2 × 2 interactions, using repeated contrasts (contrasting adjacent time points) for Time carried out via IBM SPSS Statistics for Windows, Version 28. This results in the ability to determine the adjacent time bins where transitions/change in BOLD signal dominate the significant interaction. Instead of reporting significance values for these contrasts, we report which BOLD signal adjacent timepoint changes/transitions (increases or decreases) dominate the significant Time × Condition interaction. These dominant changes are indicated in the BOLD-change figures with large brackets, complete with the dominant transition time bins printed on the top section of the brackets.

### 2.4. Tasks

Two of the tasks were designed to study working memory, specifically, WML46D04 was developed to study functional brain networks essential for verbal working memory (Sanford et al., 2020c) and the SCAP task was developed to study spatial working memory and further characterize working memory in schizophrenia (Poldrack et al., 2016). SA and PCBADE, were primarily designed to study semantic association and decision making in schizophrenia. In particular, SA was created to examine the BOLD mechanisms underlying semantic association (Woodward et al., 2015), while the PCBADE task focused on investigating evidence integration (Lavigne, Menon, Moritz, et al., 2020; Lavigne, Menon, & Woodward, 2020). RSPM, was developed as part of a broader cognitive training study, with a focus on problem-solving (Zurrin et al., 2024). The seventh task, AES, was designed to study the brain networks involved in autobiographical recall and imagination (Addis et al., 2009; Momeni et al., 2025). The eighth task, MM, aimed to explore the emergence of spontaneous thoughts during mindfulness activities (Zamani, 2023). The final task, HCP-Soc (Barch et al., 2013), was designed to study the theory of mind by probing working memory, social cognition, and visual/sensory-motor responses, respectively, in healthy individuals .

#### 2.4.1. Working Memory with Varying Cognitive Load and Delay Length (WML46D04)

The WML46D04 (working memory task with load of 4 and 6 letters and delay of 0 and 4 seconds; WML46D04) was used to assess working memory performance, and the findings were published previously (Sanford et al., 2020c) merged with a thought generation task, which is not described here. In this task, participants were presented with a sequence of four or six uppercase consonants displayed for 4s. Following a 0s or 4s delay, a single probe letter was shown for 2s. Participants were instructed to indicate whether the probe letter was part of the original sequence by pressing “yes” with their right index finger or “no” with their right middle finger. The order of letter sequences was randomized across trials to prevent pattern recognition strategies. Figure 1 illustrates the breakdown of the WML46D04 task. For the analysis presented here WML46D04

**Figure 1.**
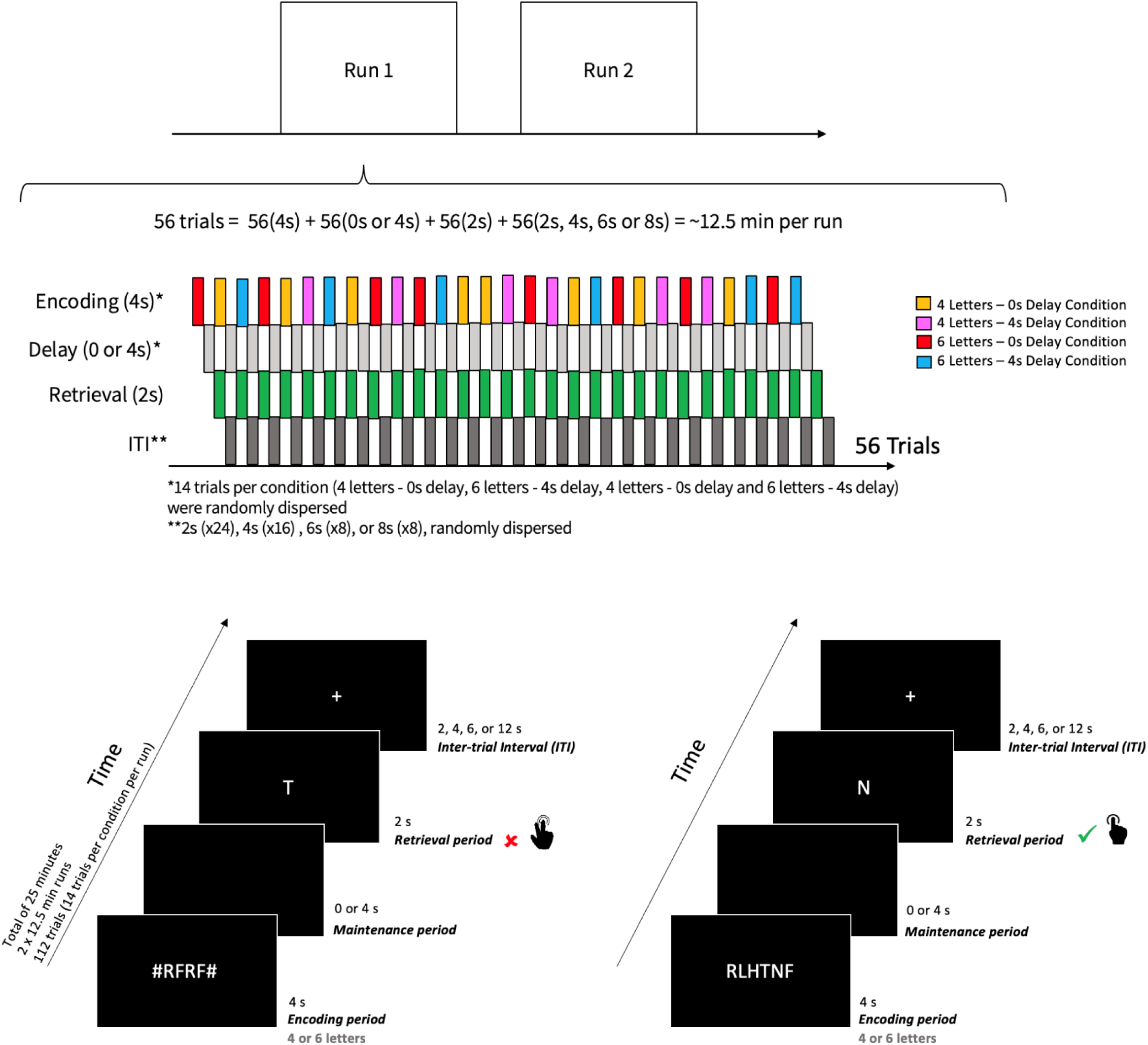
Task Diagram for Working Memory with Varying Cognitive Load and Delay Length (WML46D04).

#### 2.4.2. Spatial Capacity (SCAP)

The Spatial Capacity (SCAP) task is an item-recall task designed to assess visuospatial working memory without verbal content. The data for this task was originally collected from an OpenfMRI database (Poldrack et al., 2016) and analyzed as a part of a four-task dissertation (Sanford, 2019). In this task, illustrated in Figure 2, participants encoded and retained spatial information about target locations before making a recognition judgment. Each trial began with the presentation of a target array consisting of 1, 3, 5, or 7 yellow dots, positioned pseudo-randomly around a central fixation point. The target array remained on screen for 2 seconds, serving as the encoding period. This is followed by a delay period of either 1.5s, 3.0s, or 4.5s, during which only the central fixation point is visible. After the delay, a single green dot (the probe) appeared for 3s, and participants indicated via button press whether the probe dot was in the same position as one of the previously displayed target dots.

**Figure 2.**
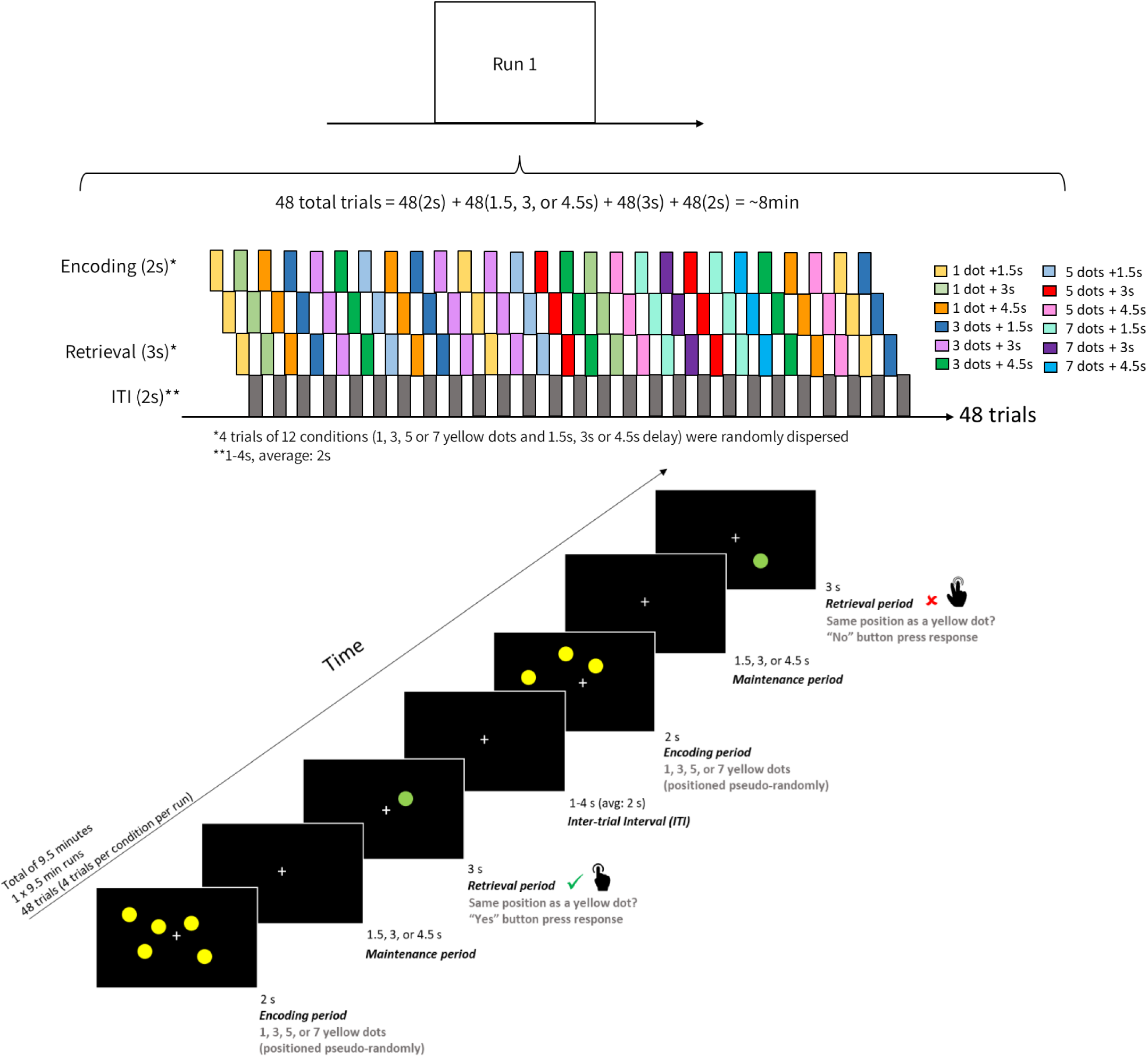
Task Diagram for Spatial Capacity (SCAP).

#### 2.4.3. Semantic Association (SA)

The Semantic Association task was initially published as a study on schizophrenia (Woodward et al., 2015). However, the current analysis focuses exclusively on results from the healthy control subjects. During this task, participants were instructed to choose a word out of three potential match options that was most closely associated with the presented prompt word that appeared in the middle of the screen. As shown in Figure 3, the prompt word was either closely related (Close condition) or distantly related (Distant condition). Participants were instructed to select the match word most closely related to the prompt word, using their index finger for the left word, middle finger for the middle word, or ring finger for the right word). If none of the options seemed related to the prompt word, they were instructed to choose the word that was most related. Each word association trial was presented for 7s, and there was a total of 90 trials. Additionally, four 9s blank trials, during which the word “Relax” was presented on the screen, were randomly inserted throughout the word association trials.

**Figure 3.**
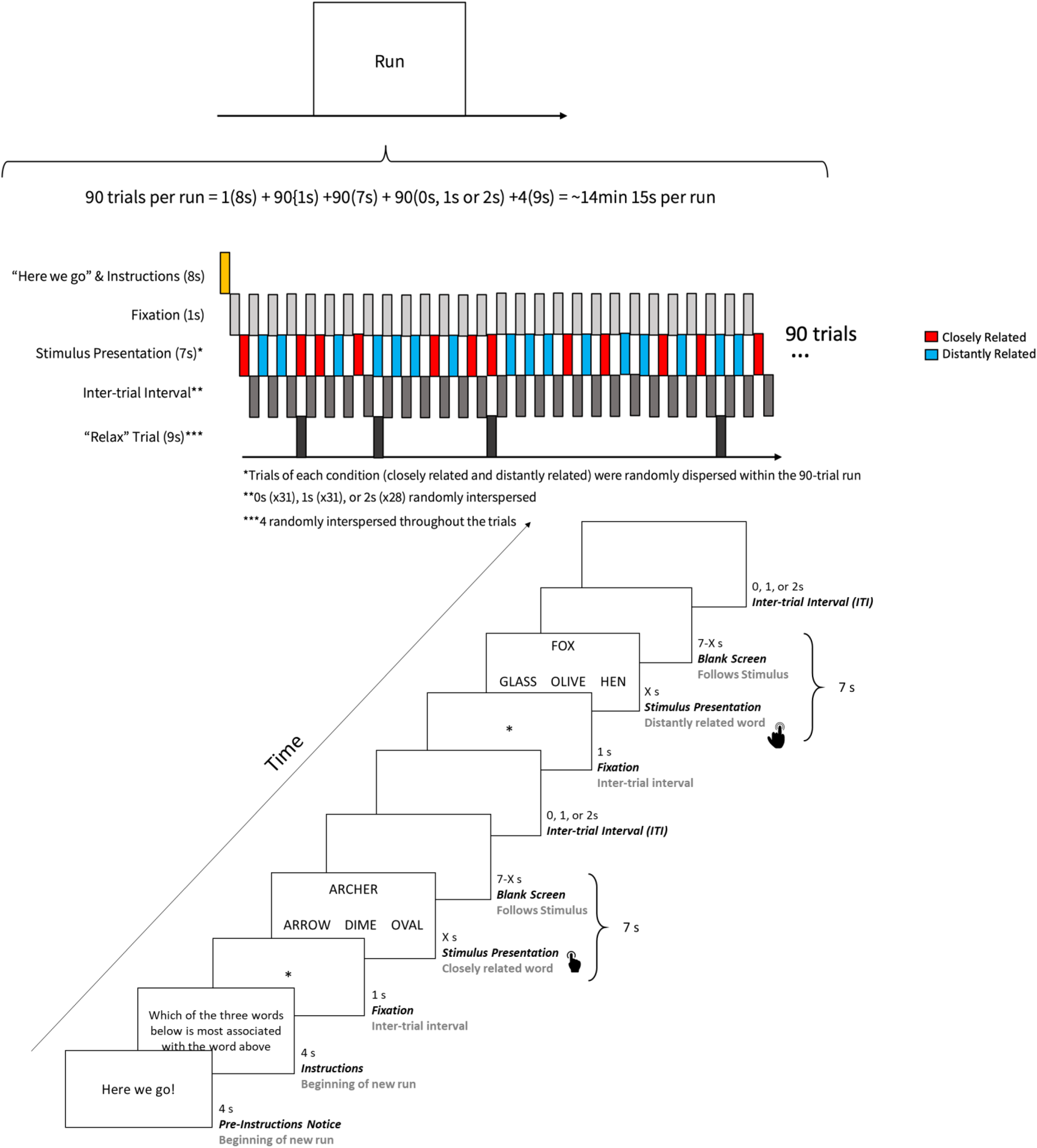
Task Diagram for Semantic Association (SA).

#### 2.4.4. Picture Bias Against Disconfirmatory Evidence (PCBADE)

The PCBADE task was developed to study evidence integration (Lavigne, Menon, Moritz, et al., 2020; Lavigne, Menon, & Woodward, 2020). Participants were presented with a partial line drawing of common objects, food, or animals, and were asked to rate whether they believed the full picture was of a word listed below the image using a dichotomous (yes/no) response interface. Following a response, they were presented with a second partial image showing more of the full picture and were asked to re-rate. Finally, the full picture was presented, which did not require a response. Figure 3 shows the timing of the experimental paradigm using a disconfirm trial with the picture “bat” and lure word “umbrella”. Conditions were based on expected response patterns (two responses per trial): Yes-Yes (YY), No-No (NN), No-Yes (NY), and Yes-No (YN). YY and NN are “confirm” conditions, since the initial rating is supported by the second rating, whereas NY and YN are “disconfirm” conditions, since the second rating contradicts the first rating.

**Figure 3.**
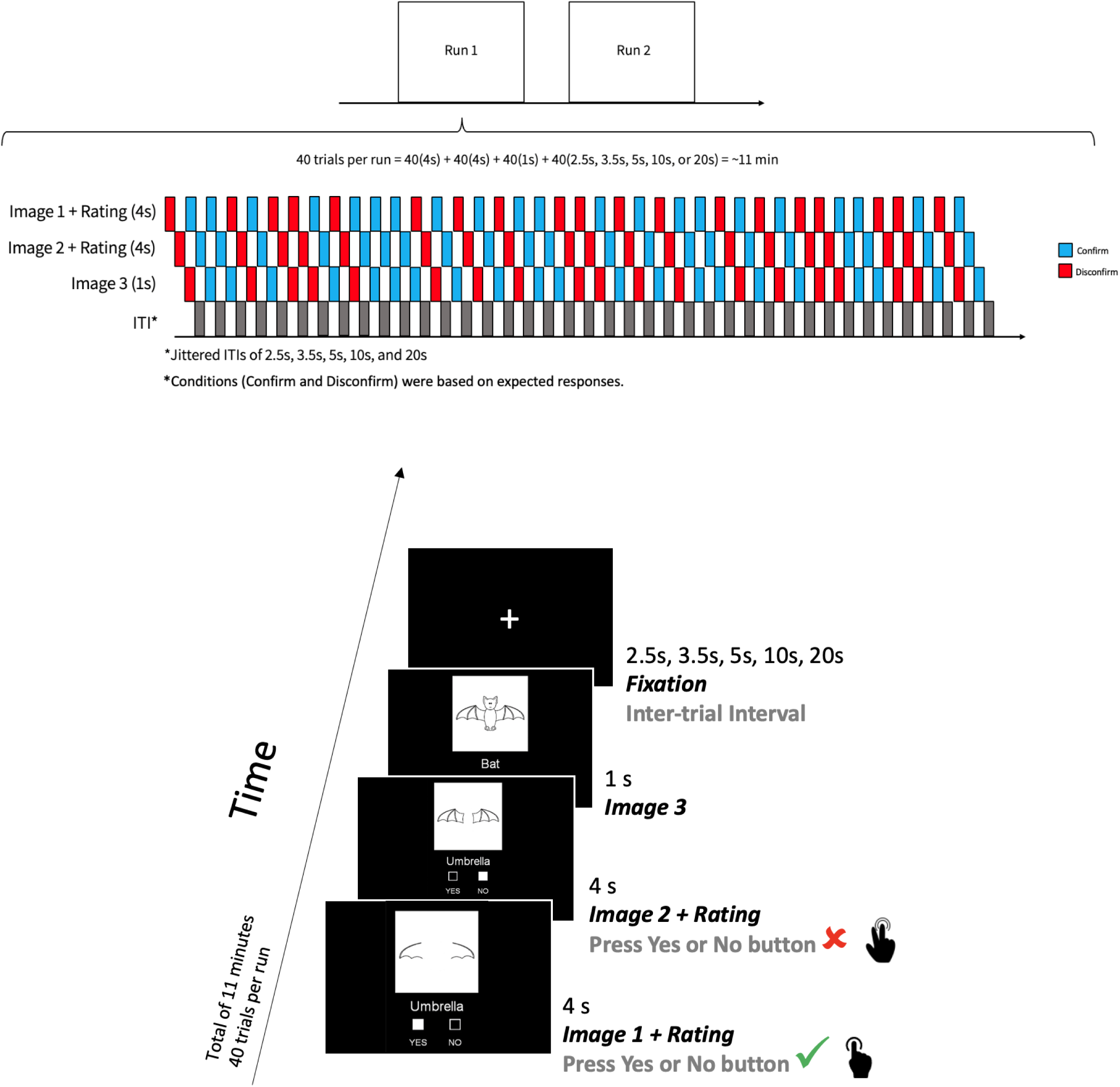
Task Diagram for Picture Bias Against Disconfirmatory Evidence (PCBADE).

#### 2.4.5. Raven’s Standard Progressive Matrices (RSPM)

The RSPM task was developed to investigate problem-solving and cognitive training, and details about RSPM are available in published work (Zurrin et al., 2024). Shown in Figure 4 is a timeline of one block of stimulus presentations in the RSPM task. A matrix was presented for 5s, after which there was an 8s response period whereby an answer was highlighted in red, and the participants responded whether the highlighted answer was correct or incorrect via a button press with their index or middle finger, respectively. One block consisted of ten matrix problems. The block shown (Figure 4) displays medium difficulty RSPM problems. After participants finished reading the instructions, 15s of fixation were presented at the beginning of each block. One second of fixation was presented before each 13s trial. The task concluded with 15s of fixation. One easy block, one medium block, and one hard block were presented to each participant.

**Figure 4.**
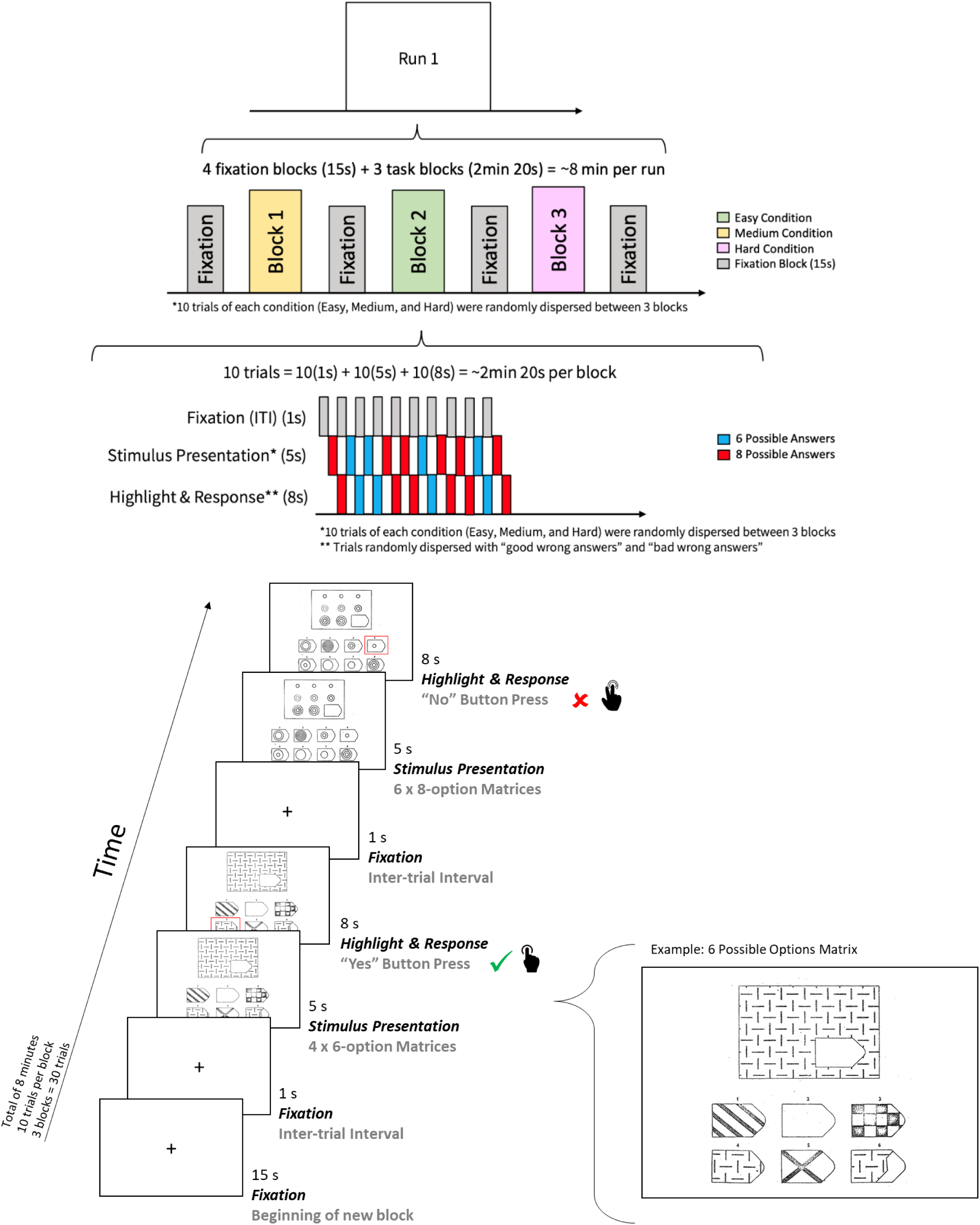
Task Diagram for Raven’s Standard Progressive Matrices (RSPM).

#### 2.4.6. Autobiographical Event Simulation (AES)

The AES task was developed to study autobiographical recall and imagination. Details of the AES task have been previously published (Addis et al., 2009) and a summary is presented here. The AES task (Figure 5) was comprised of four different conditions: (1) imagining past autobiographical events (Imagine-Past), (2) imagining future autobiographical events (Imagine-Future), (3) recalling past autobiographical events (Recall), and (4) a semantic association control task (Semantic Association). In the Imagine-Past and Imagine-Future trials, the construction phase involved using the specified details (i.e., person, place, and object) on the cueing slide to imagine a temporally and contextually specific event that could have occurred in the past 5 years but did not, or that might occur in the next 5 years, respectively. In the Recall condition, construction involved participants remembering memories corresponding to each of the specified details on the cueing slide. Once participants had imagined or recalled events, they pressed a button on the response box which marked the end of the construction phase and the beginning of the elaboration phase. During the elaboration phase, participants were required to elaborate and expand on the events by imagining or recalling as much details as possible from a field perspective (i.e., seeing the event from the perspective of being there). Immediately after elaboration, two rating scales were presented, each for 5s. The first one was a five-point scale assessing the amount of detail retrieved or imagined (1 = vague with no/few details; 5 = vivid and highly detailed). The second one was a binary scale assessing whether the event imagined or retrieved was experienced primarily from a field or observer perspective (1 = saw the event through my own eyes; 5 = saw myself from an external perspective).

**Figure 5.**
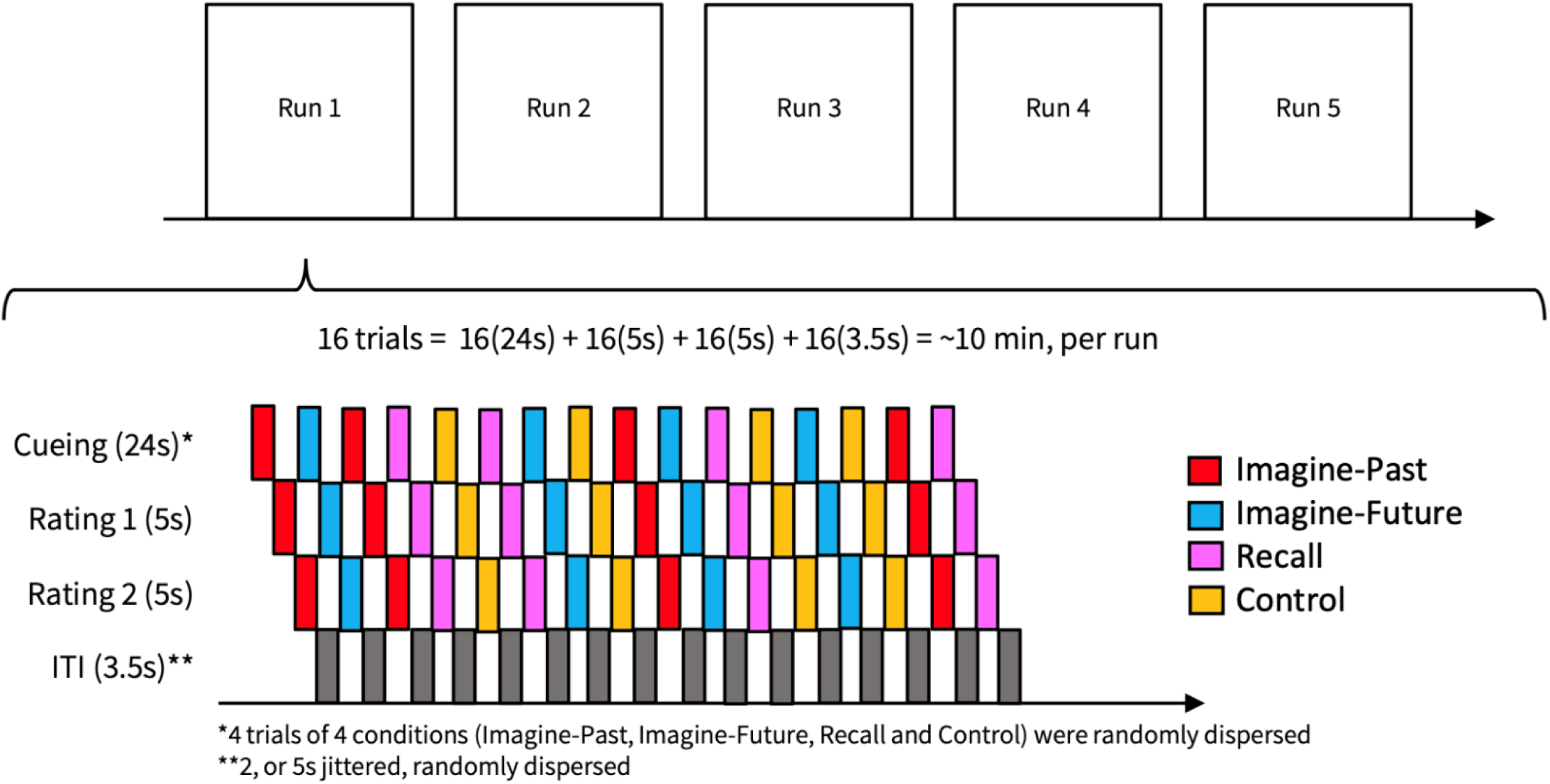
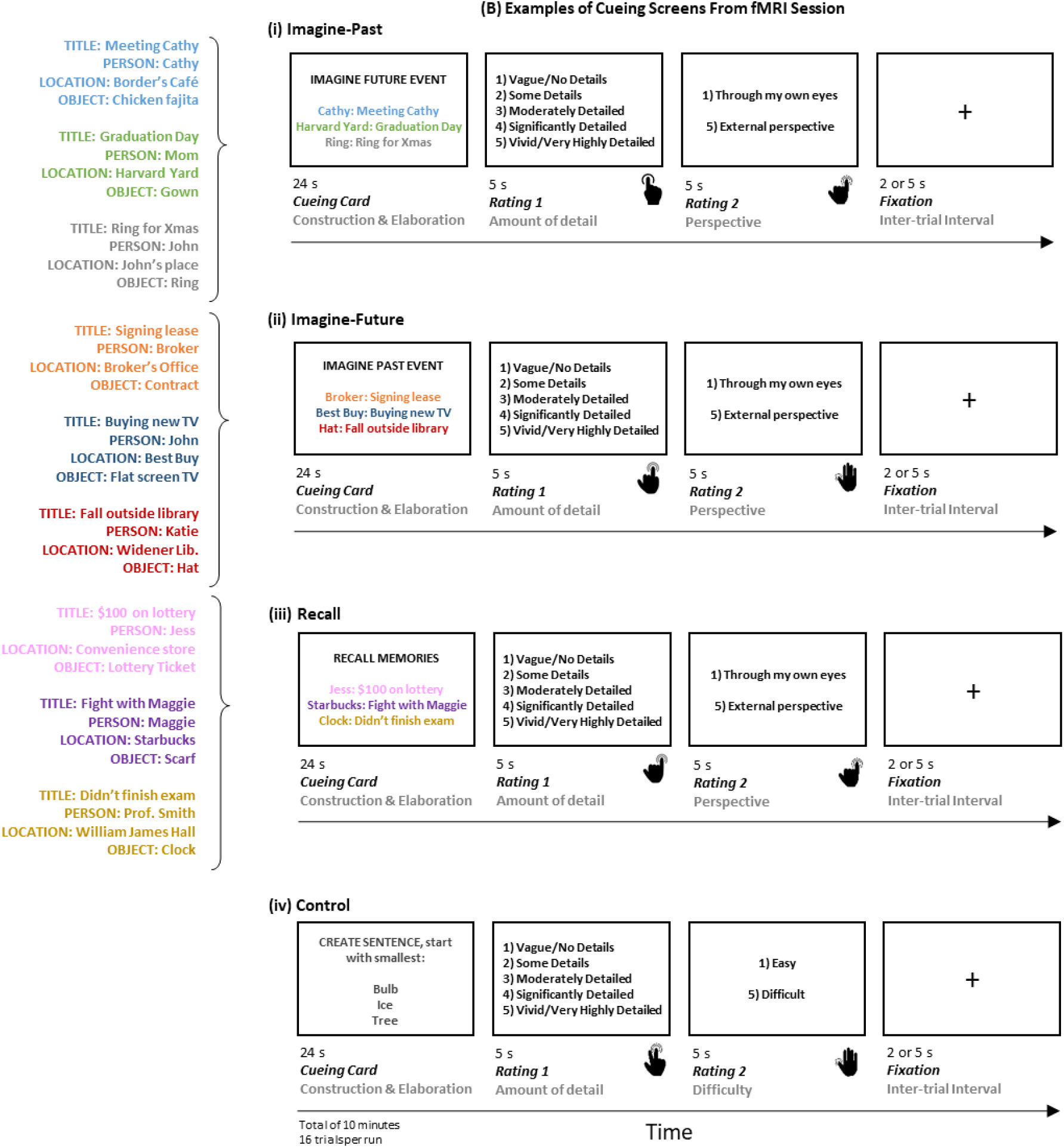
Task Diagram for Autobiographical Event Simulation (AES).

**Figure 6.**
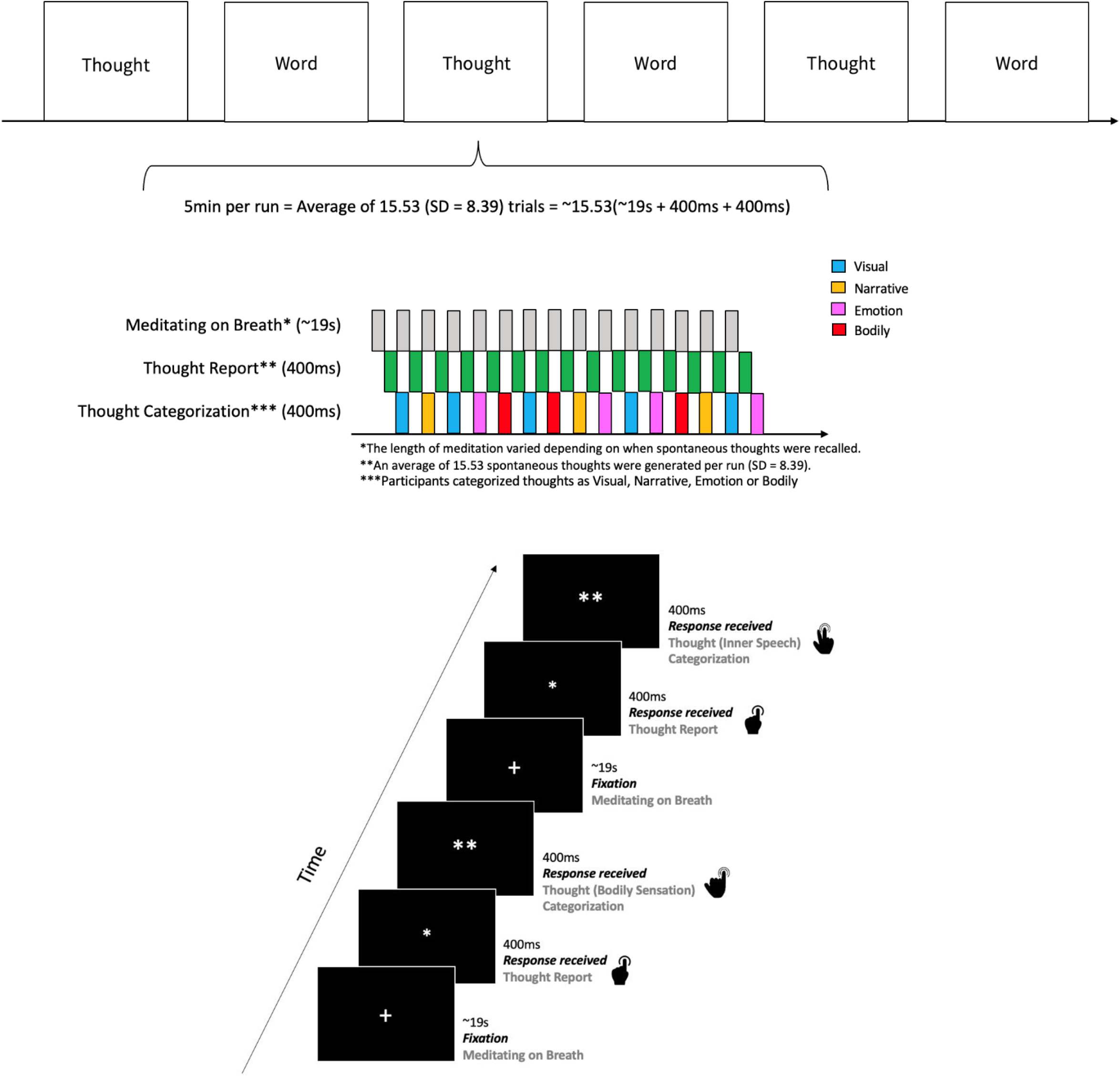
Task Diagram for Mindfulness Meditation (MM).

Similar to the AES trials, each semantic association control trial began with a 24s construction and elaboration phase. During the construction phase, participants were required to order the three objects by physical size and insert them into the following sentence: “X is smaller than Y is smaller than Z”. Once participants had silently said the sentence to themselves, they made a button-press, marking the end of the construction phase. Participants then elaborated on the representation of the nouns, generating as many details about their meaning and physical attributes as possible. Following the elaboration phase, two rating scales were presented for 5s each. The first one was a five-point scale assessing the amount of detail generated during the elaboration phase (1 = vague with no/few details; 5 = vivid and highly detailed). The second one was a binary scale assessing task difficulty (1 = easy; 5 = difficult).

#### 2.4.7. Mindfulness Meditation (MM)

The MM task design presented here was modified from a previously published manuscript by (Ellamil et al., 2016) and was a follow-up analysis based on data collected for a publicly available masters’ thesis (Zamani, 2023). The experimental procedure for this study was designed to investigate the cognitive mechanisms underlying spontaneous thoughts using a within-subjects yoked-control design. The task consisted of two alternating 5-minute conditions: a Thought condition followed by a Word condition, both structured to mirror key aspects of mindfulness meditation. The underlying task design remained the same across both conditions: participants focused on their breath while gazing at a white fixation cross on a black screen (see Figure 15). During the Thought condition, they were instructed to press a button (left pointer finger) as soon as they noticed an internally generated thought arise, then they were required to press a second button to categorize the thought’s modality (right pointer finger = visual, right middle finger = narrative/inner speech, right ring finger = emotion, right pinky finger = bodily sensation). In the Word condition, participants viewed a word presented at random on-screen and responded in the same manner, categorizing the word’s referential content rather than its mode of presentation. In both conditions, a single asterisk (*) was displayed after each first button press (400ms), and a double asterisk (**) after each second button press (400ms), providing feedback on response completion. Participants were instructed to return attention to their breath following each response. Only the Thought condition is presented here.

#### 2.4.8. Human Connectome Project – Social (HCP-Soc)

The analysis of the 500 subject HCP-Soc task has not been published. However, this analysis is from the publicly available Human Connectome Project data set (Barch et al., 2013). In the HCP-Soc Human Connectome social task, participants were shown a short video clip (20s each) featuring different objects, such as squares, circles, triangles. Example images taken from these video clips are depicted in Figure 7. During each video (Castelli et al., 2000; Wheatley et al., 2007), the objects either interacted in a specific manner (Mental Condition Block) or moved randomly (Random Condition Block). After watching the video, participants had to choose one of three options via a button press: (1) the objects had a social interaction (i.e. the shapes seemed to consider each other’s feelings or thoughts), (2) not sure, or (3) no interaction (i.e. movement appeared random). Each video block was followed by a 15s fixation block. The task consisted of a total of 5 Mental and 5 Random blocks, with fixation blocks interwoven between each video block.

**Figure 7.**
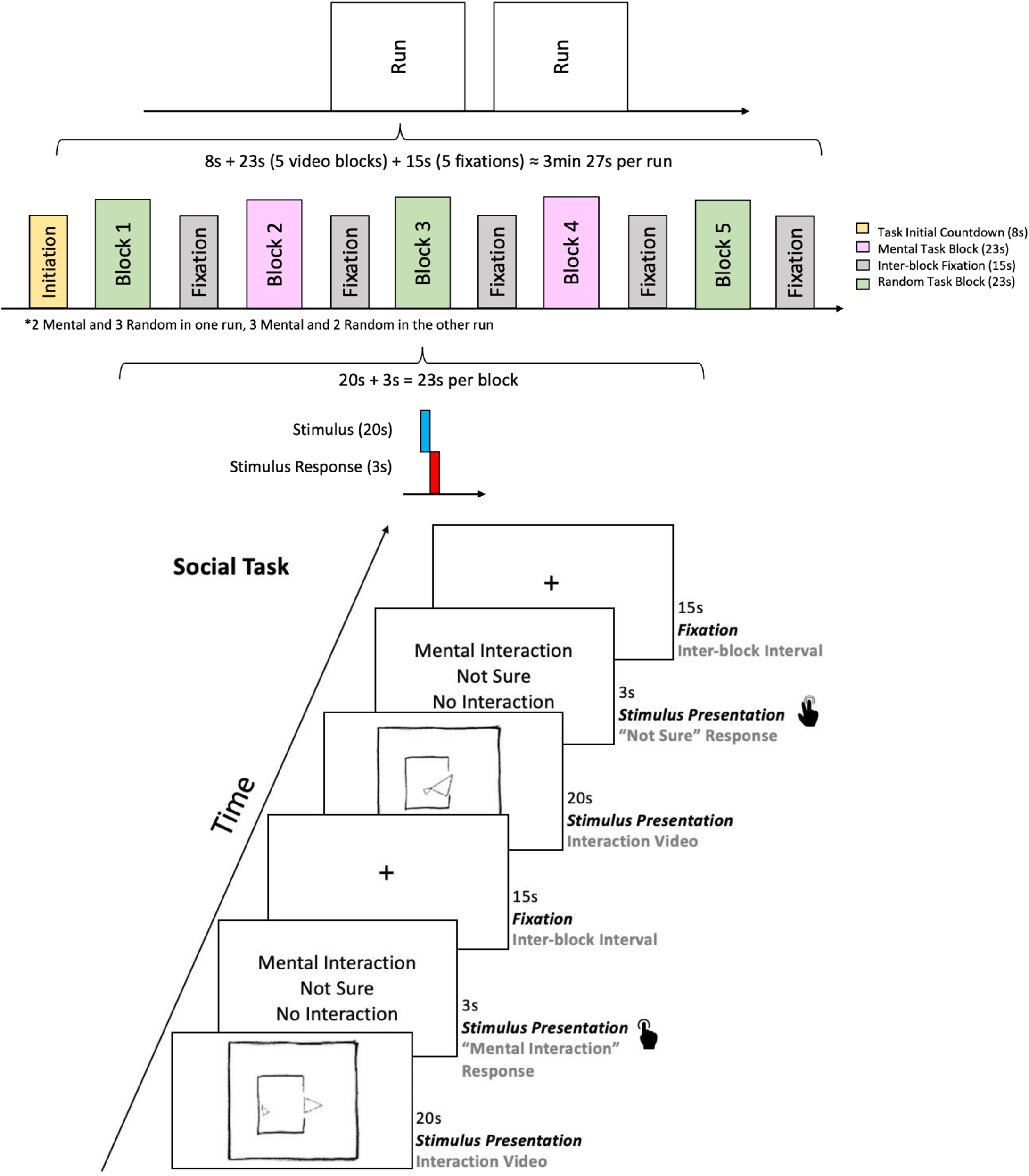
Task Diagram for Human Connectome Project – Social (HCP-Soc).

### 2.5. Subjects

A total of 753 subjects were included in these analyses, all of whom were screened to ensure the absence of mental health diagnoses and brain disorders. Demographic information for each study is shown in Table 1. For the published studies using the same samples reported here (Lavigne, Menon, Moritz, et al., 2020; Sanford, 2019; Sanford et al., 2020c; Woodward et al., 2015; Zurrin et al., 2024) further details are provided in the original publications. The WML46D04-TG and AES tasks involved only healthy controls and were two of the 10 task images that were averaged to create the 2020 anatomical template images. In particular, 27 of the 32 participants who had completed the TG task had scans that were averaged twice to make the original DMB template.

**Table 1.** DMB participant characteristics for each study are reported here. The % subject overlap with anatomical template image indicates that the N subjects of the study task comprised the listed percentage of the total subjects included in the creation of the anatomical template image. The % scan set overlap with anatomical template image indicates that the N subjects of the study task contributed the listed percentage of the scan sets to the scan sets included in the creation of the anatomical template image. The % image overlap with anatomical template indicates that the study’s image, with specified classification *Z* score, comprised the listed percentage of the total anatomical images included in the creation of the anatomical template image.

| Study | Task | N | Age | Handedness (R:L:M) | Sex (M:F) | % scan set overlap with template | % image overlap with template | Classification Z score |
| --- | --- | --- | --- | --- | --- | --- | --- | --- |
| WML46D 04-TGT | WML46D 04 | 37 | 28.16 (8.70) | 32:3:2 | 10:27 | 6.06% | 10% | 1.77 |
|  | TGT | 32 | 28.75 (8.58) | 29:3:0 | 19:13 | 5.24% |  |  |
| SCAP | SCAP | 44 | 31.61 (8.84) | 44:0:0 | 20:24 | 0% | 0% | 0.92 |
| SA | SA | 30 | 26.80 (6.11) | 28:2:0 | 13:17 | 0% | 0% | 0.77 |
| PCBADE | PCBADE | 41 | 35.44 (12.59) | 39:2:0 | 19:22 | 0% | 0% | 1.09 |
| RSPM | RSPM | 56 | 31.10 (6.10) | 56:0:0 | 23:33 | 0% | 0% | 0.80 |
| AES | AES | 17 | 22.00 (2.66) | 17:0:0 | 9:8 | 2.78% | 10% | 0.91 |
| MM | MM | 18 | 55.72 (13.64) | 14:4:0 | 6:11* | 0% | 0% | 0.81 |
| HCP-Soc | HCP-Soc | 500 | 26.5 (2.37) | † | 223:277 | 0% | 0% | 0.91 |

\*One participant's sex was unreported. † Handedness information missing

The SCAP, SA, PCBADE, RSPM, MM and HCP-Soc images presented here provide entirely independent replications of the DMB pattern-based anatomy as all participant scan sets and the resultant anatomical images were not in any way included in, nor were overlapping with, the original 2020 anatomical template image (Percival et al., 2020). All study procedures were fully approved by the Research Ethics Board of the corresponding location where the study took place. All participants provided informed consent before taking part in the study.

## 3. Results

### 3.1. Anatomical Patterns

The anatomical patterns that are characteristic of the DMB cognitive mode were nick-named as follows: (1) Snowman Nose, (2) Prominent Medial Temporal Dots, (3) In Flight, (4) Tripod, and (5) Angel Wings. These nick-names are based on five visually identifiable patterns which, when occurring as a group, regardless of which other patterns are present in positive or negative loadings, are put forward as indicative of the presence of DMB. That is to say, the presence of the Snowman Nose + Prominent Medial Temporal Dots + In Flight + Tripod + Angel Wings in a to-be-identified brain image, regardless of other patterns present in the brain image, indicates the presence of DMB, and only DMB. In Table 2, we present examples of each pattern emphasized in the DMB cognitive mode patterns first made public in 2020 (Percival et al., 2020). It is important to emphasize that none of these patterns appear in their pure form on any other cognitive mode images (Percival, Chen, & Woodward, 2024). We use image thresholds as a microscope-style tool to search for these patterns and therefore do not use strict cutoff values (Chen & Woodward, 2024). In addition to Table 2, in Supplementary Table S1 the X Y Z MNI coordinate peaks on each slice are depicted, overlaid on 7-network Yeo/Buckner/Choi template (Buckner et al., 2011; Choi et al., 2012; Yeo et al., 2011), and also provided are the Harvard Oxford, Automated Anatomical Labeling (AAL), Brodmann Region, and Yeo/Buckner/Choi network number for cognitive mode activation peaks.

**Table 2.**
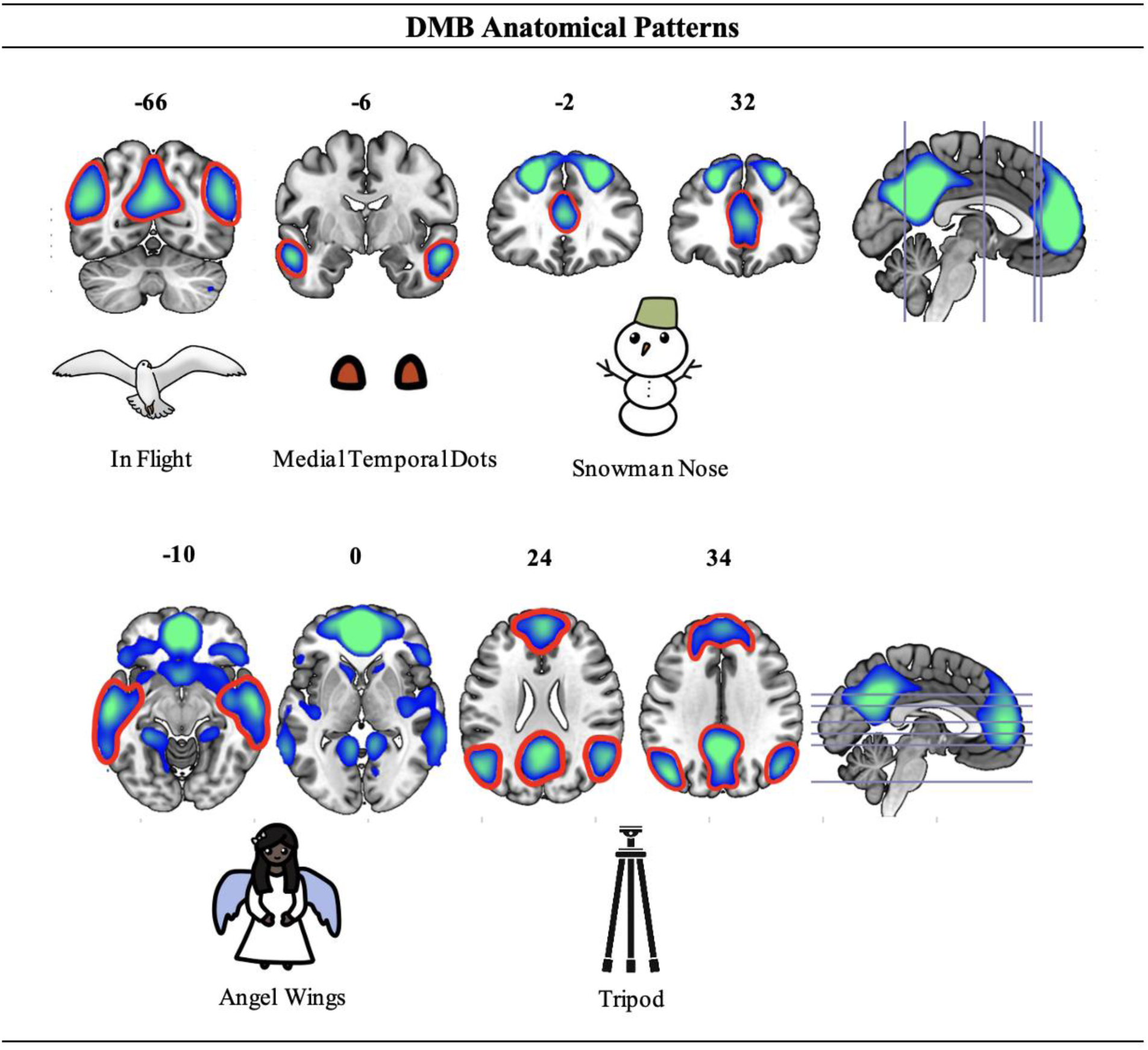
Default Mode B (DMB) Anatomical Patterns. MNI slice numbers are displayed for coronal and axial views, in neurological convention (left is left). Blue/green represents negative component loadings and regions of deactivation. Red lines and circles highlight key patterns that characterize DMB: (1) Snowman Nose (coronal slice 32 and 37), (2) Prominent Medial Temporal Dots (coronal slices -6 and -2), (3) In Flight (coronal slice -66), (4) Tripod (axial slice 24), and (5) Angel Wings (axial slices -10 and 0). None of these five occur in their pure form, in isolation or in co-occurrence, in any other cognitive mode (Percival, Chen, & Woodward, 2024).

As the DMB is a task-negative network, we focus on patterns of task-induced deactivations instead of activations. Snowman Nose, an anatomical pattern specific to DMB, emerges from deactivations in the anterior cingulate gyrus (ACC) and the bilateral superior frontal gyrus (SFG) on coronal slice 32 in Table 2 (Percival, Chen, & Woodward, 2024). In this pattern, deactivations in the SFG resemble the snowman’s eyes, while the deactivation in the ACC forms the snowman’s nose. Moving posteriorly, on coronal slices -6 and -2, pronounced bilateral deactivations in the middle temporal gyrus (MTG) give rise to the Prominent Medial Temporal Dots pattern. The In Flight pattern is observed on coronal slice -66, where distinctive wing-shaped deactivations are found in the bilateral lateral occipital cortex (LOC). On the same slice, deactivations in the precuneus (PCUN) may be visualized as resembling the head and body of a bird in flight. Distinct DMB patterns also appear on axial slices. As depicted in axial slice 24, the Tripod pattern emerges from four deactivation clusters in the SFG, PCUN, and LOC, which can be connected in a way that resembles the structure of a tripod. Lastly, deactivations on axial slices -10 and 0 mark the Angel Wings pattern. On axial slice -10, deactivation in the medial frontal gyrus (MFG) corresponds to the angel’s head, while deactivations in the subcallosal cortex (SCC) and posterior MTG correspond to the angel’s body and wings, respectively. More superiorly, on axial slice 0, the angel’s wingtips may be represented by deactivations in the temporooccipital part of the MTG, whereas the angel’s feet may be visualized by deactivations in the lingual gyrus (LING).

#### 3.1.1. Working Memory with Varying Cognitive Load and Delay Length - Thought Generation Merge (WML46D04-TG)

The fMRI-CPCA results presented here are published as part of a seven-cognitive mode set of results (Sanford et al., 2020c). The anatomical patterns that are characteristic of DMB were observed in the merged analysis of the WML46D04 and TG tasks. The *Z* score for DMB was 1.77, which corresponds to a Pearson’s *r*^2^ value of .89. As shown in Table 3, we can see that In Flight, Medial Temporal Dots, Snowman Nose, Angel Wings, and Tripod were all very well retrieved.

**Table 3.**
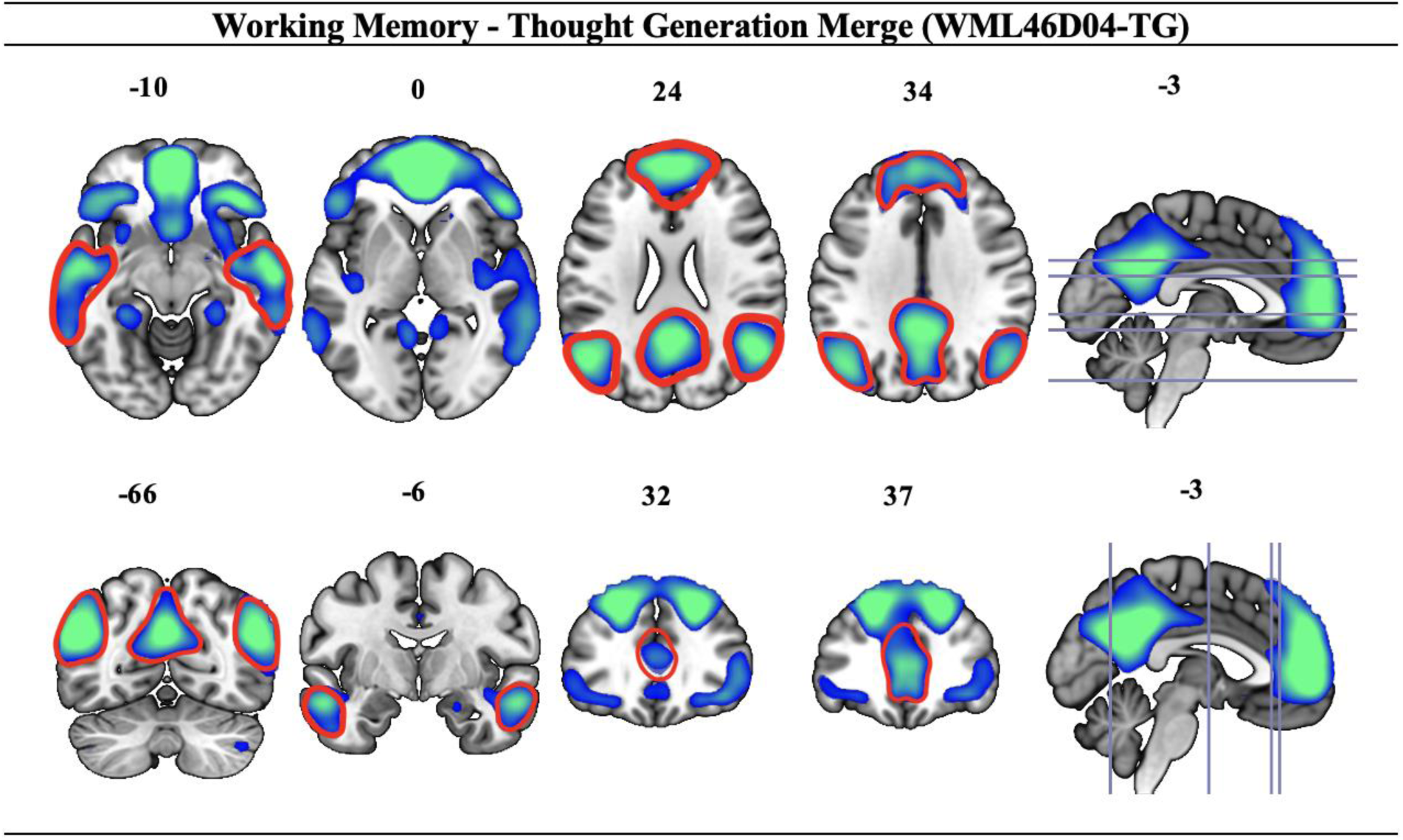
Default Mode B (DMB) anatomical patterns for Working Memory with Vary Cognitive Load and Delay Length - Thought Generation Merge (WML46D04-TG). Red lines indicate the presence of the DMB anatomical patterns *Z* = 1.77. This image was not included in the original 2020 DMB anatomical template image.

#### 3.1.2. Spatial Capacity (SCAP)

The fMRI-CPCA results presented here are part of a published dissertation two cognitive mode set of results (Sanford, 2019). The anatomical patterns that are characteristic of DMB were observed in the SCAP task. The *Z* score for DMB was 0.92, which corresponds to a Pearson’s *r*^2^ value of .53. As shown in Table 4, we can see that In Flight, Medial Temporal Dots, Snowman Nose, Angel Wings, and Tripod were all well retrieved.

**Table 4.**
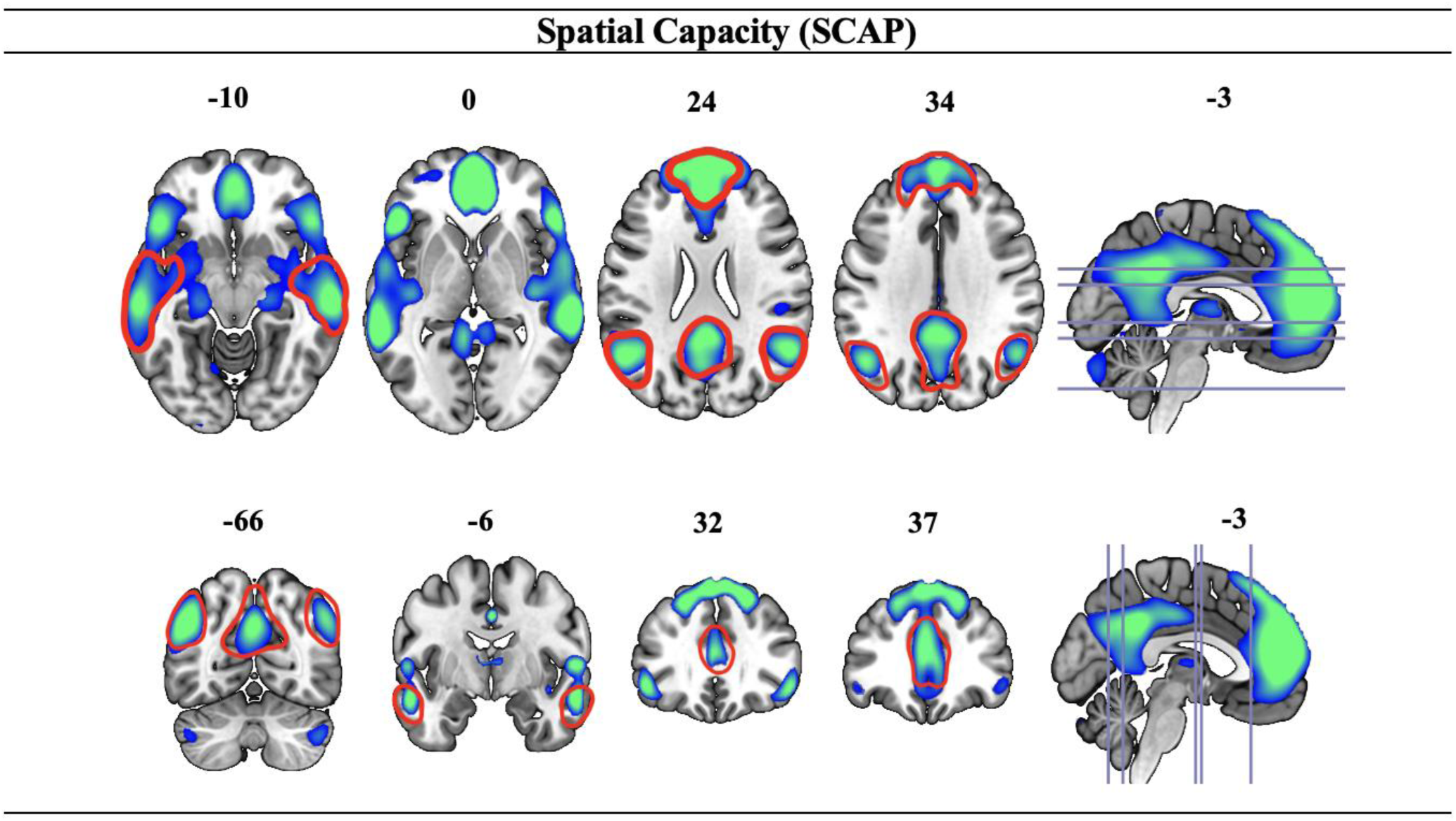
Default Mode B (DMB) anatomical patterns for Spatial Capacity (SCAP). Red lines indicate the presence of the DMB anatomical patterns *Z* = 0.92. This image was not included in the original 2020 DMB anatomical template image.

#### 3.1.3. Semantic Association (SA)

The fMRI-CPCA results presented here are a part of a five-cognitive mode set of results, with a similar set including both schizophrenia and healthy participants (Woodward et al., 2015). The *Z* score for DMB was 0.77, corresponding to a Pearson’s *r^2^* value of .42. As depicted in Table 5, Angel Wings, Medial Temporal Dots and Snowman are formed. However, In Flight was poorly formed, and Tripod was missing the top pattern.

**Table 5.**
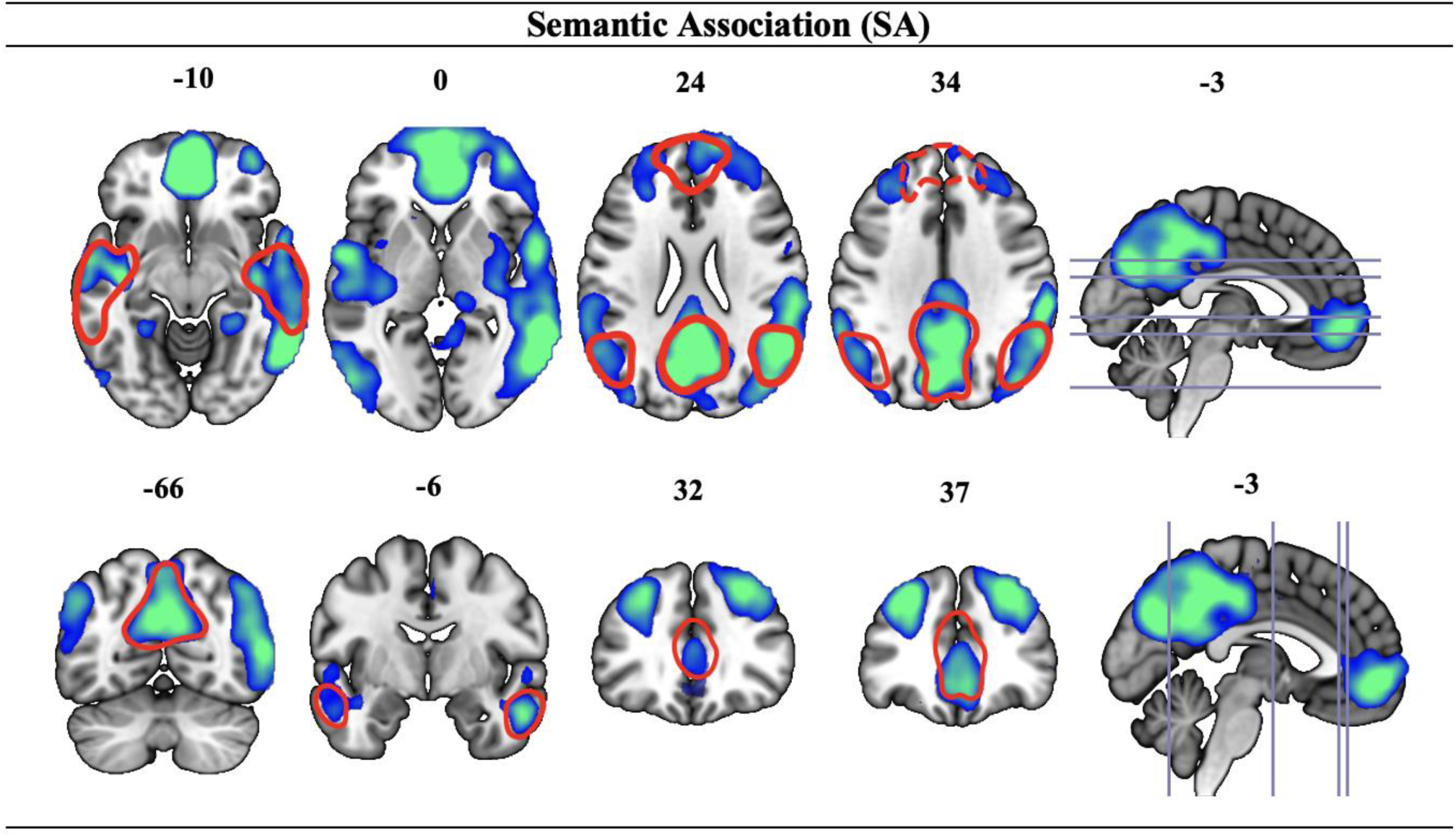
Default Mode B (DMB) anatomical patterns for Semantic Association (SA). Red lines indicate the presence of the DMB anatomical patterns *Z* = 1.77. This image was not included in the original 2020 DMB anatomical template image.

#### 3.1.4. Picture Completion Bias Against Disconfirmatory Evidence (PCBADE)

The fMRI-CPCA results presented here for PCBADE are published as part of a three-cognitive mode set of results (Lavigne, Menon, Moritz, et al., 2020). The *Z* score for DMB was 1.09, which corresponds to a Pearson’s *r^2^* value of .64. As depicted in Table 6, the DMB anatomical patterns Angel Wings, In Flight, Medial Temporal Dots, and Snowman were all present. The top head piece of the Tripod pattern was not observed.

**Table 6.**
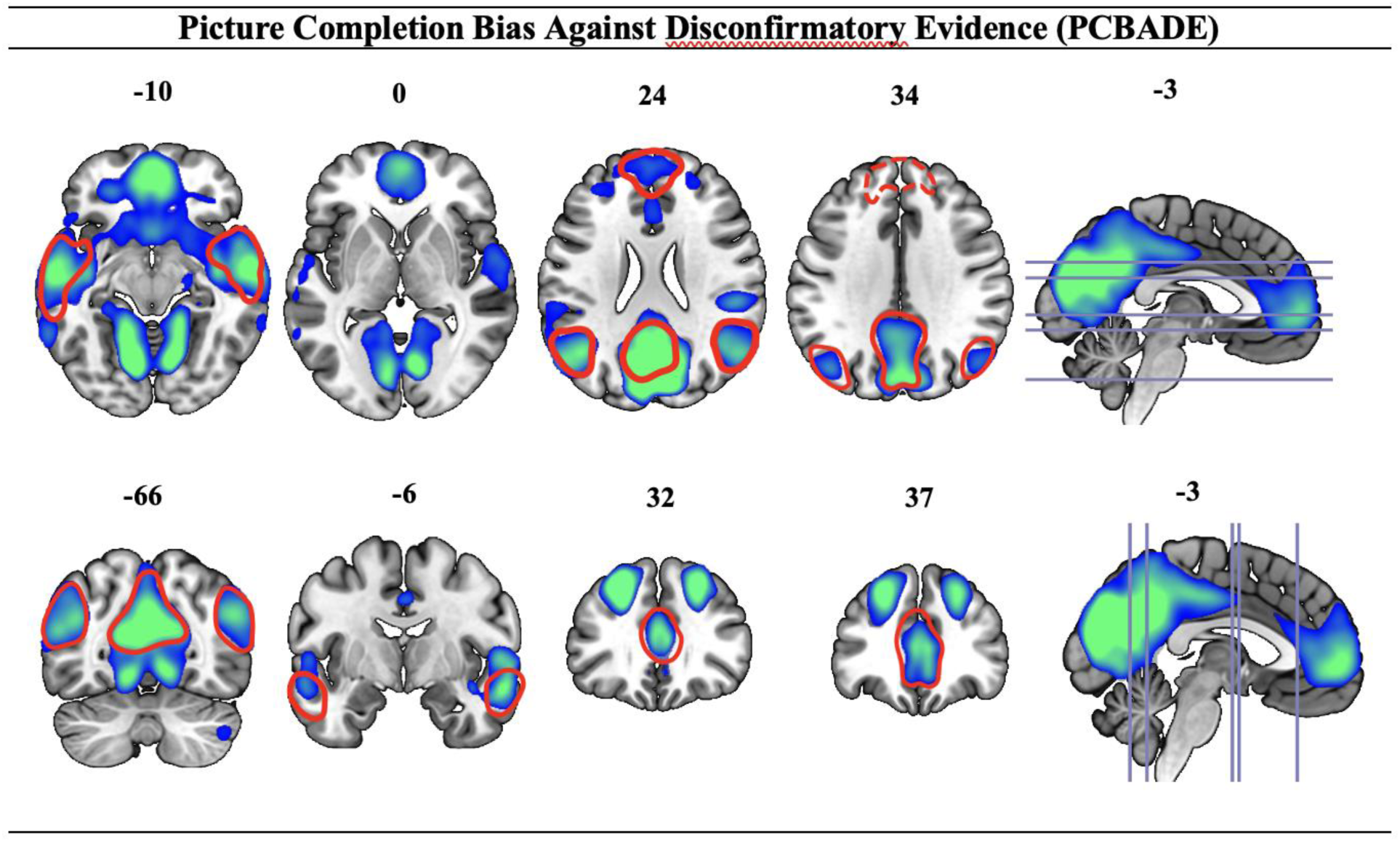
Default Mode B (DMB) patterns for Picture Completion Bias Against Disconfirmatory Evidence (PCBADE). Red lines indicate the presence of the DMB anatomical patterns *Z* = 1.09. This image was not included in the original 2020 DMB anatomical template image.

#### 3.1.5. Raven’s Standard Progressive Matrices (RSPM)

The fMRI-CPCA results presented here for RSPM are published as part of a five-cognitive mode set of results (Zurrin et al., 2024). This RSPM image (Table 7) was completely independent from the original 2020 DMB anatomical template image. The anatomical patterns that are characteristic of DMB were observed. The *Z* score for DMB was 0.80, which corresponds to a Pearson’s *r*^2^ value of .44. In Table 7 we see that DMB patterns Angel Wings, Tripod, In Flight, Medial Temporal Dots, and Snowman were all present.

**Table 7.**
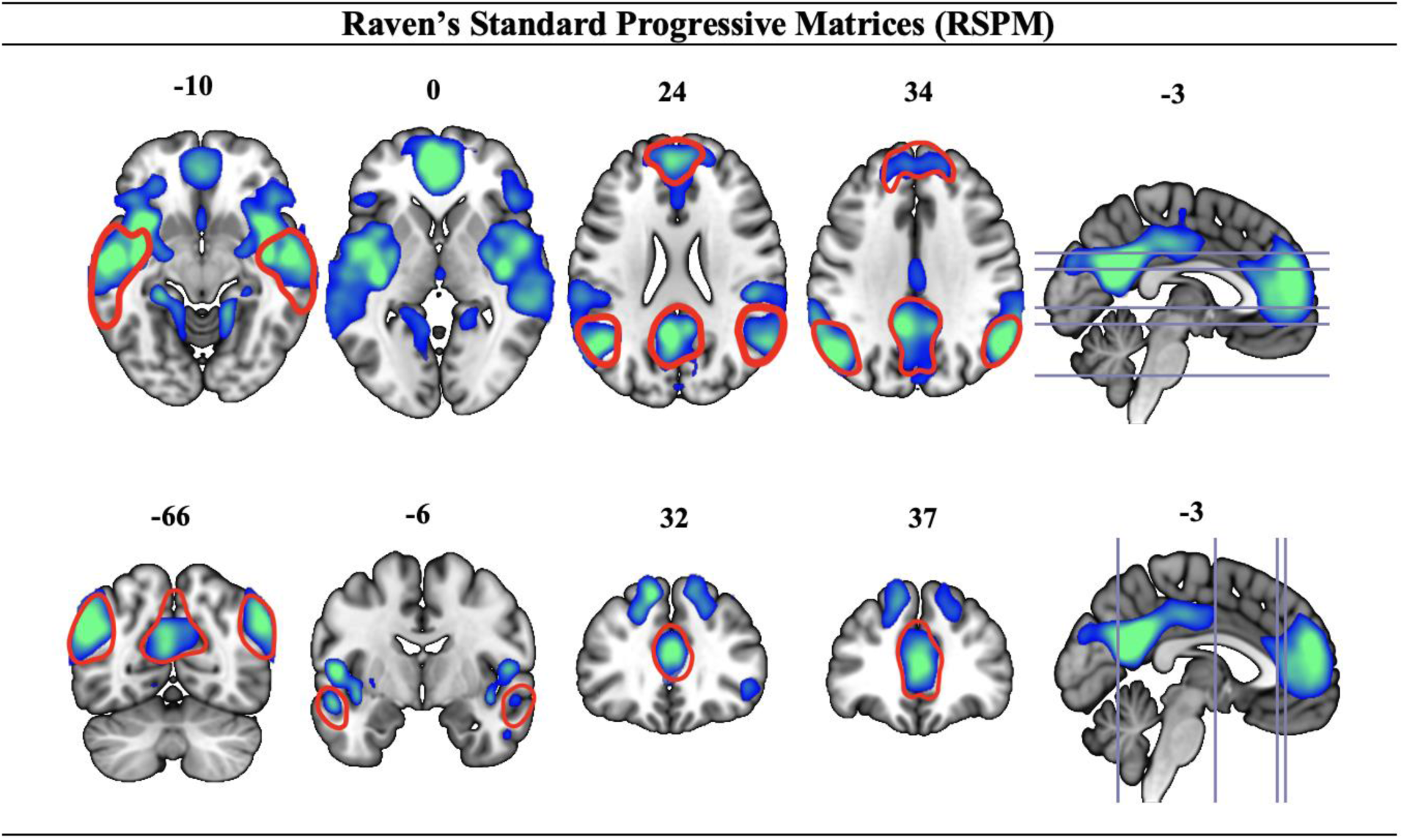
Default Mode B (DMB) anatomical patterns for Raven’s Standard Progressive Matrices (RSPM). Red lines indicate the presence of the DMB anatomical patterns Task. *Z* = 0.80. This image was not included in the original 2020 DMB anatomical template image.

#### 3.1.6. Autobiographical Event Simulation (AES)

The AES fMRI-CPCA results presented here are published as part of a five-cognitive mode set of results (Momeni et al., 2025). The *Z* score for DMB was 0.91, corresponding to a Pearson’s *r*^2^ value of .52. Table 8 shows that In Flight, Medial Temporal Dots, Snowman Nose, Angel Wings, and Tripod were all clearly present.

**Table 8.**
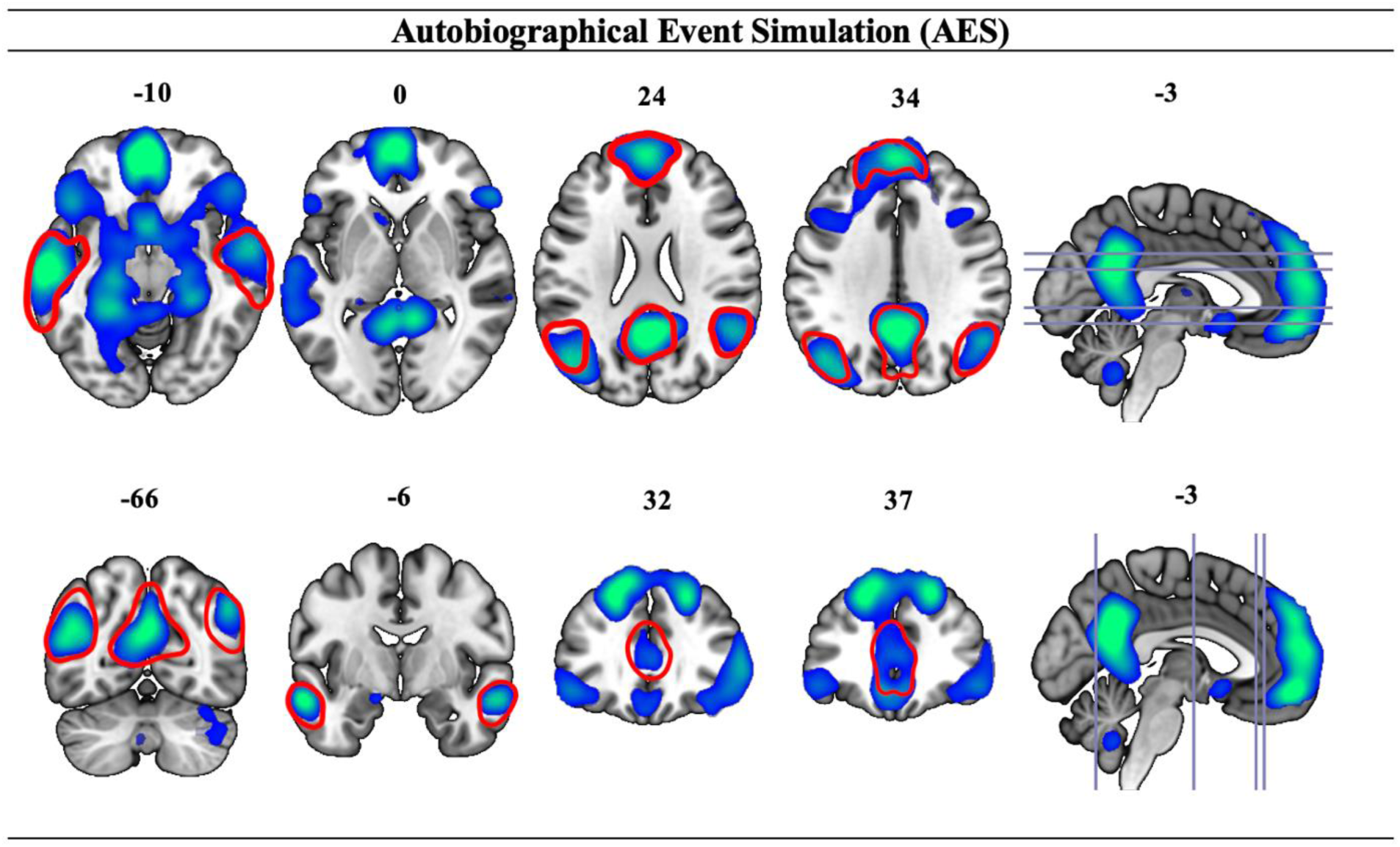
The Default Mode B (DMB) anatomical patterns for Autobiographical Event Simulation (AES). Red lines indicate the presence of the DMB anatomical patterns *Z* = 0.91. This image was not included in the original 2020 DMB exemplar image.

#### 3.1.7. Mindfulness Meditation (MM)

The MM fMRI-CPCA results presented here are a follow-up analysis on other work (Zamani, 2023), and are part of an unpublished four-cognitive mode set of results. The *Z* score for DMB was 0.81, corresponding to a Pearson’s *r*^2^ value of .45. The Angel Wings, Tripod, In Flight, and Medial Temporal Dots were all clearly present in Table 9 The Snowman Nose was also present but was poorly defined relative to prototypical DMB pattern.

**Table 9.**
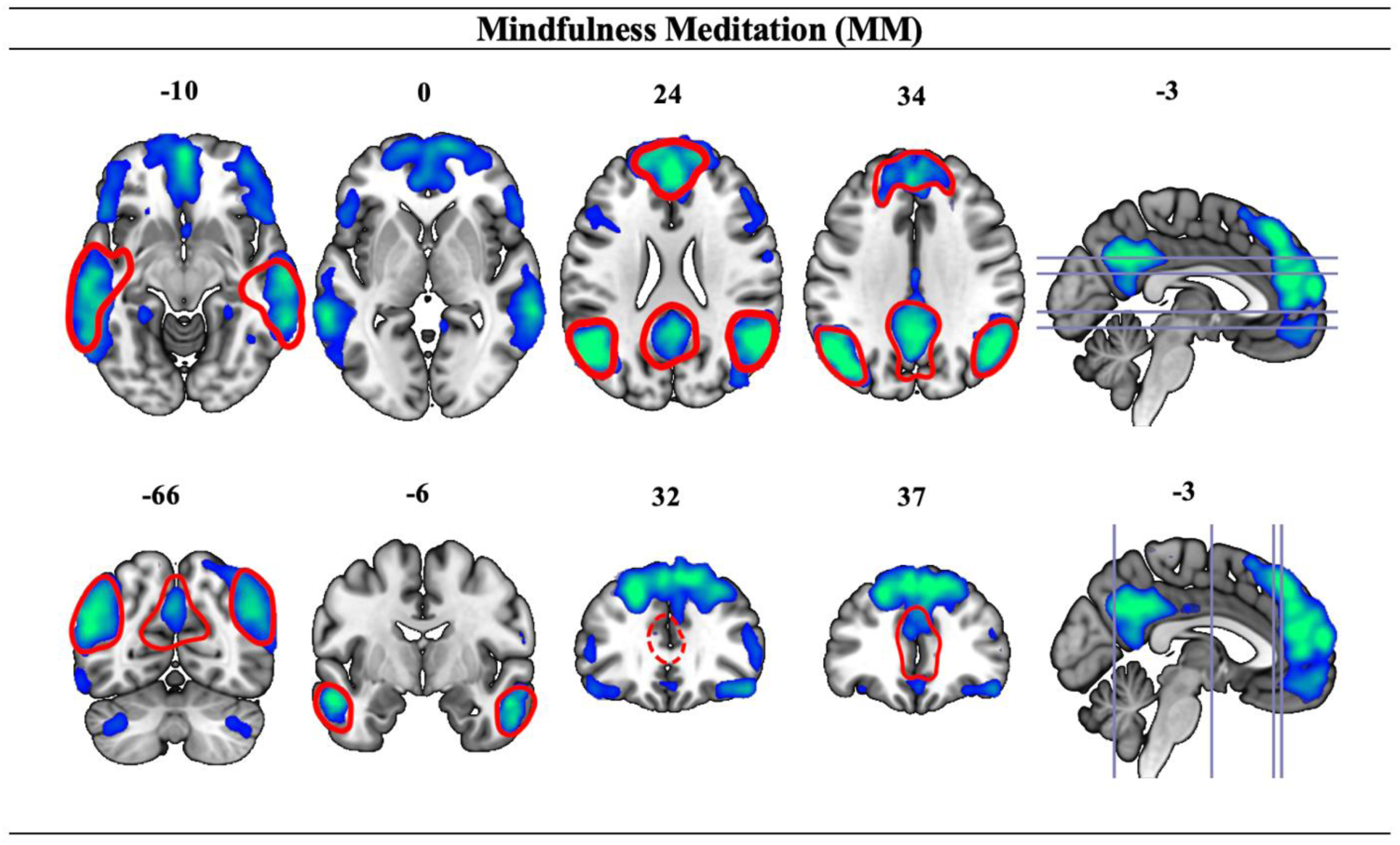
The Default Mode B (DMB) anatomical patterns for Mindfulness Meditation (MM). Red lines indicate the presence of the DMB anatomical patterns *Z* = 0.81. This image was not included in the original 2020 DMB exemplar image.

#### 3.1.8. Human Connectome Project – Social (HCP-Soc)

The fMRI-CPCA results presented here for HCP-Soc are part of an unpublished eight-cognitive mode set of results, which has not been published but belongs to an analysis of the publicly available Human Connectome Project dataset (Barch et al., 2013). The *Z* score for the DMB network was 0.91, corresponding to Pearson’s *r*^2^ value of 0.52. As shown in Table 10, the DMB patterns Angel Wings, Tripod, and Snowman are all well-replicated. The right dot of Medial Temporal Dots was not observed.

**Table 10.**
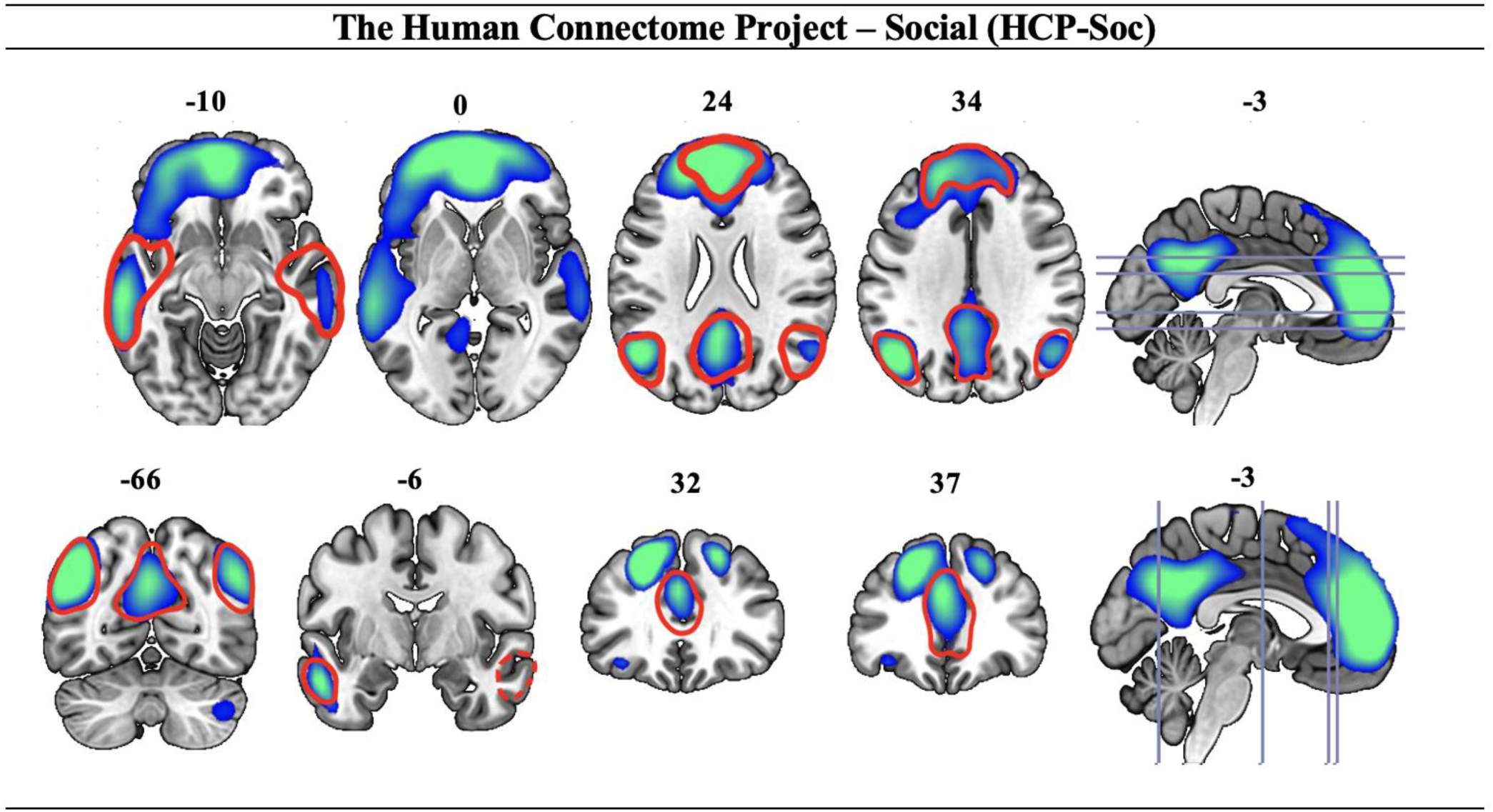
The Default Mode B (DMB) anatomical patterns for Human Connectome Project - Social (HCP-Soc). Red lines indicate the presence of the DMB anatomical patterns *Z* = 91. This image was not included the original 2020 DMB anatomical template image.

### 3.2. Temporal Patterns

#### 3.2.1. Working Memory with Varying Cognitive Load and Delay Length - Thought Generation Merge (WML46D04-TG)

For the WML46D04 task, using a 2 (Load - 4, 6 Letters) × 2 (Delay - NoDelay, 4sDelay) × 10 (Time) RM-ANOVA on DMB revealed significant main effects of Time, *F*(9,324) = 38.57, *p* < .001, *η_p_^2^* = .52, Load *F*(1,36) = 22.26, *p* < .001, *η_p_ ^2^* = .38, and Delay *F*(1,36) = 24.83, *p* < .001, *η_p_ ^2^*= .41. Additionally, there was a significant Load × Time interaction, *F*(9,324) = 6.29, *p* < .001, *η_p_^2^* = .15, a significant Delay × Time interaction, *F*(9,324) = 16.37, *p* < .001, *η_p_^2^* = .31, and a significant Load × Delay interaction, *F*(1,36) = 7.36, *p* < .05, *η_p_^2^* = 0.17. The Delay × Time interaction was dominated by the BOLD changes between 11s-13s (see Figure 8), whereby the 0-Delay conditions (solid lines) were decreasing activity due to being post-peak, while the 4-Delay conditions (dotted lines) were still increasing activity due to later peaks, demonstrating extended and delayed peaks corresponding to increased memory delay. Load dependence is also clear from observation of Figure 8 (red lines > blue lines).

**Figure 8.**
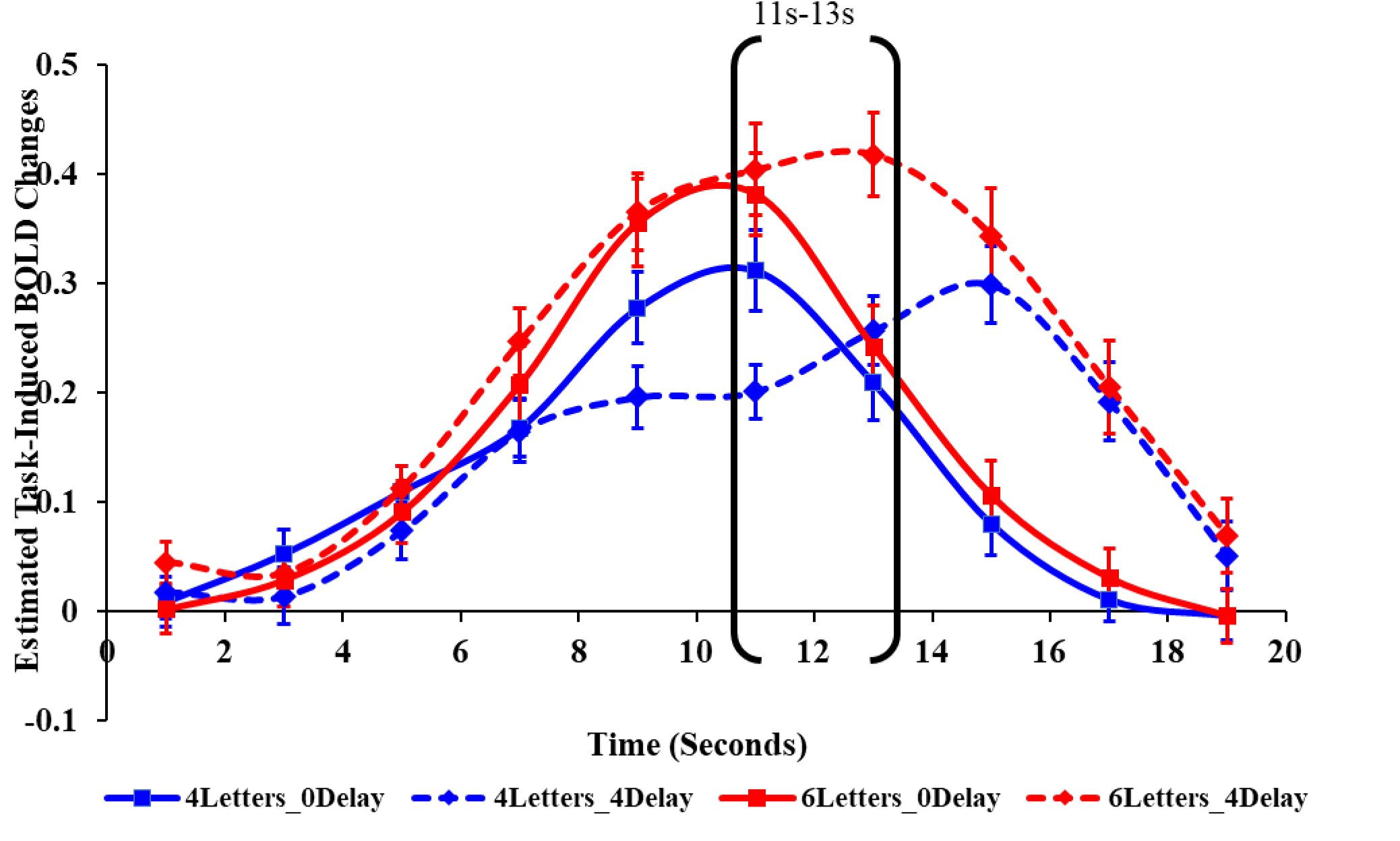
Estimated task-induced BOLD signal changes for WML46D04. Mean finite impulse response (FIR)-based predictor weights plotted as a function of post-stimulus time. Positive predictor weights plotted here reflect DMB deactivation, corresponding to the blue/green voxels in the associated anatomical patterns. Brackets emphasize the task-induced BOLD changes dominating the Delay (NoDelay, 4sDelay) × Time interaction, whereby the 0-Delay conditions (solid lines) were decreasing activity due to being post-peak, while the 4-Delay conditions (dotted lines) were still increasing activity due to later peaks. The fMRI repetition time (TR) for this study was 2s.

For the TG task, using a 2 (Condition - Hearing, Generating) × 10 (Time) RM-ANOVA for DMB indicated a significant main effect of Time, *F*(9,279) = 15.62, *p* < .001, *η_p_^2^* = 0.34.

However, there were no significant main effect or interaction involving Condition (both *p*s > .30).

#### ***4.2.2*** Spatial Capacity (SCAP)

By using a 2 (Load-5-Dots, 7-Dots) × 3 (Delay-1.5s, 3.0s, 4.5s) × 10 (Time) RM-ANOVA for the SCAP task, all main effects and interactions were significant (all *p*s < .01), with effect sizes *η_p_^2^* ranging from .03 - .52. Here we present the results from Load × Time, *F*(9,990) = 3.81, *p* < .001, *η_p_^2^* = .03 (see Figure 9), and Delay × Time *F*(18,1980) = 9,44, *p* < .001, *η_p_^2^* = .08 (see Figure 10) with the interactions caused by load- and delay-dependent effects in the expected directions. The Load × Time interaction was dominated by adjacent contrasts of 5-7s and 13-15s due to increasing deactivation with load (see Figure 9). The Load × Delay interaction was dominated by adjacent contrasts of 9-13s due to increasing duration of deactivation with delay period (see Figure 10).

**Figure 9.**
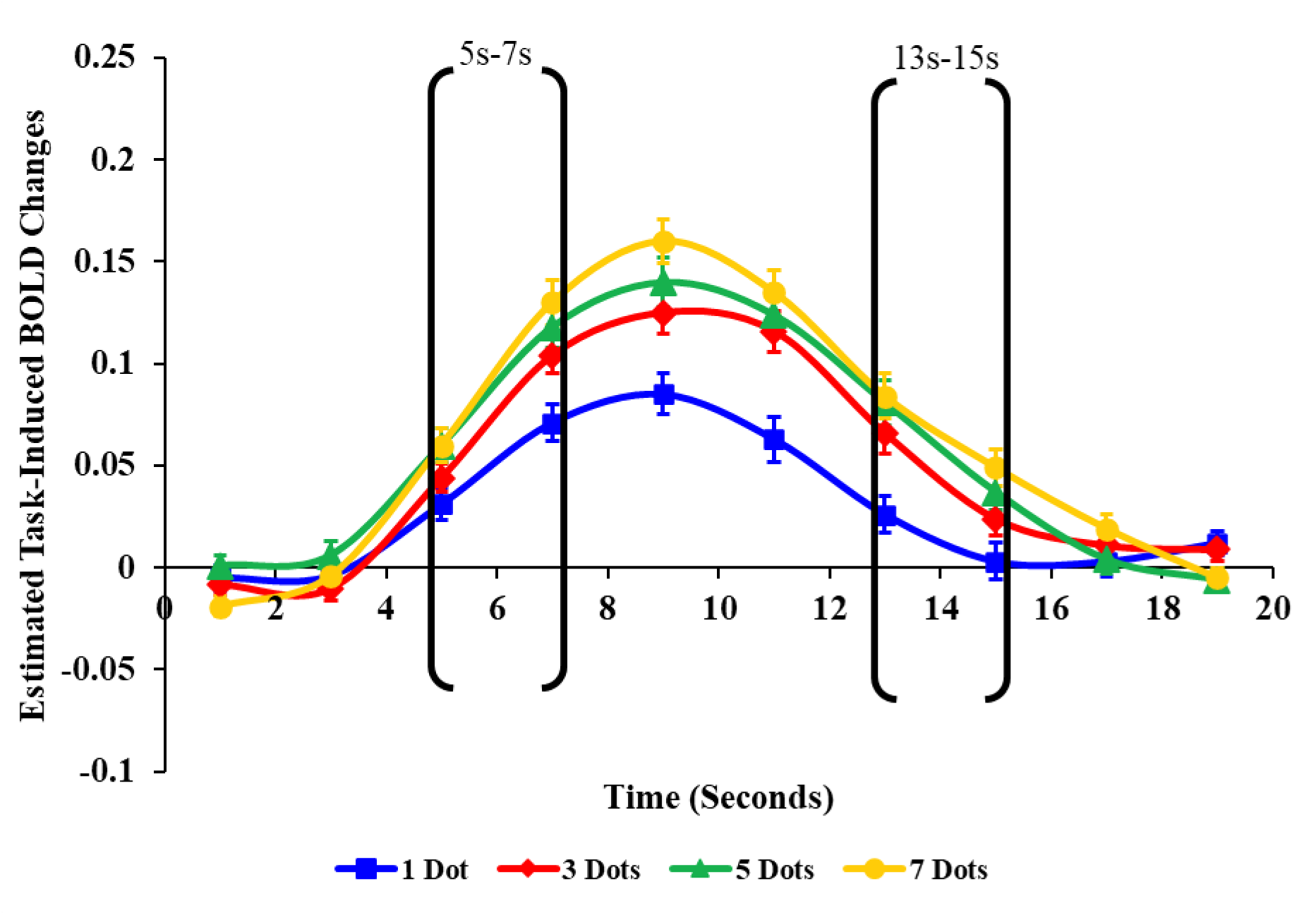
Estimated task-induced BOLD signal changes for SCAP. Mean finite impulse response (FIR)-based predictor weights plotted as a function of post-stimulus time. Positive predictor weights plotted here reflect DMB deactivation, corresponding to the blue/green voxels in the associated anatomical patterns. Brackets emphasize the task-induced BOLD changes dominating the Cognitive Load (1, 3, 5 and 7) × Time interaction, due to increasing deactivation with increasing load. The fMRI repetition time (TR) for this study was 2s.

**Figure 10.**
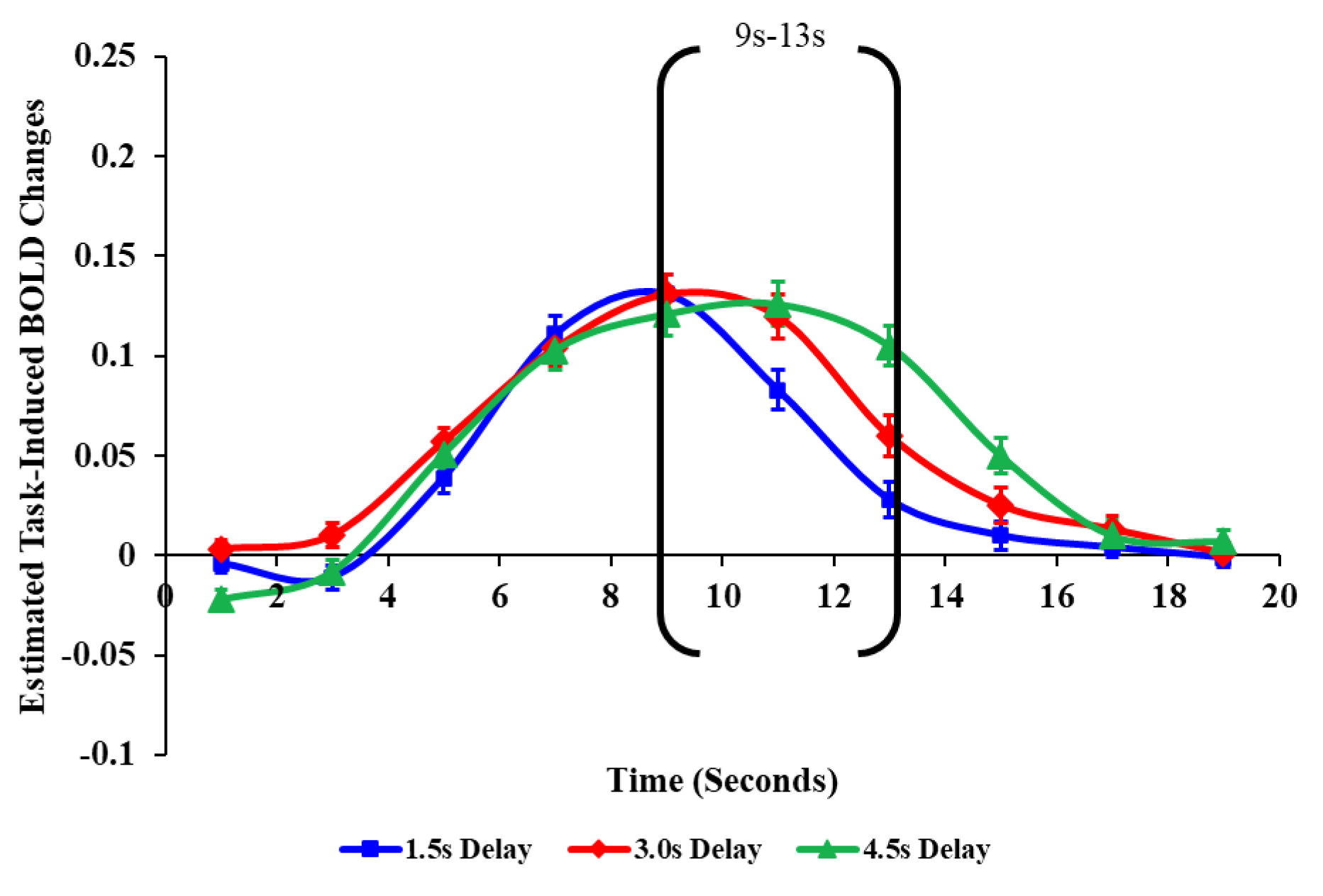
Estimated task-induced BOLD signal changes for SCAP. Mean finite impulse response (FIR)-based predictor weights plotted as a function of post-stimulus time. Positive predictor weights plotted here reflect DMB deactivation, corresponding to the blue/green voxels in the associated anatomical patterns. Brackets emphasize the task-induced BOLD changes dominating the Delay (1.5s, 3s, 4.5s) × Time interaction, due to the 1.5 delay condition deactivations peaking earlier and then decreasing as the 3.0 delay condition peaks. The fMRI repetition time (TR) for this study was 2s.

#### 4.2.3. Semantic Association (SA)

For the SA task, a 2 (Relatedness - Close, Distant) × 10 (Time) RM-ANOVA analysis indicated a significant main effect for Relatedness, *F*(1,29) = 31.18, *p* < .001, *η_p_ ^2^*= .52. Also present was a significant Relatedness × Time interaction, *F*(9,261) = 6.05, *p* < .001, *η_p_^2^*= .17. This interaction was dominated by contrasts 5s to 7s and contrasts 15s to 17s. From 5s to 7s, the Distant condition exhibited a higher increase to peak deactivation compared to the Close condition, resulting in a higher deactivation peak for the Distant condition at approximately 9s (see Figure 11). Since the Distant condition was more strongly deactivated, between 15s and 17s, its deactivation decreased more steeply compared to the Close condition to return to the baseline level.

**Figure 11.**
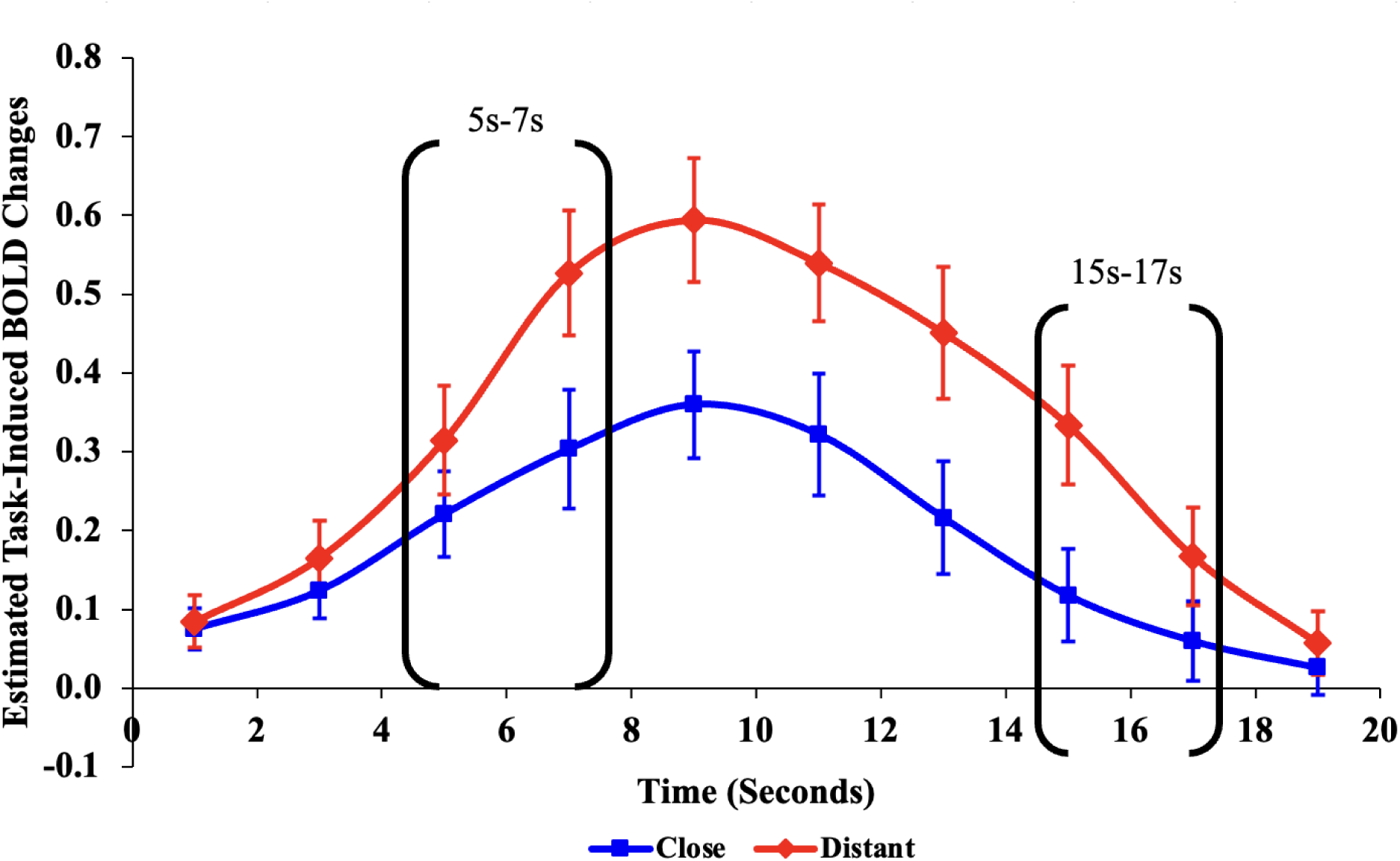
Estimated task-induced BOLD signal changes for SA. Mean finite impulse response (FIR)-based predictor weights plotted as a function of Relatedness and Time. Positive predictor weights plotted here reflect DMB deactivation, corresponding to the blue/green voxels in the associated anatomical patterns. Brackets indicate the task-induced BOLD changes dominating the Relatedness × Time interaction due to greater DMB deactivation in the Distant condition. The fMRI repetition time (TR) for this study was 2s.

#### 4.2.3. Picture Bias Against Disconfirmatory Evidence (PCBADE)

By using a 2 (Evidence Type - Confirm, Disconfirm) × 2 (Evidence Order - YesFirst, NoFirst) × 10 (Time) RM-ANOVA for the PCBADE task, significant main effects were observed for Time, *F*(9,351) = 73.01, *p* < .001, *η_p_^2^* = .65, and a significant Evidence Type × Time interaction, *F*(9,351) = 3.57, *p* < .001, *η_p_^2^ =* .84, but other main effects and interactions were not significant (all *p*s > .05). The Evidence Type × Time interaction was dominated by contrasts between time bins 13s to 15s, due to the steeper peak deactivation for the Disconfirm condition (Figure 12, red line), returning closer to baseline more rapidly than the Confirm condition (Figure 12, blue line).

**Figure 12.**
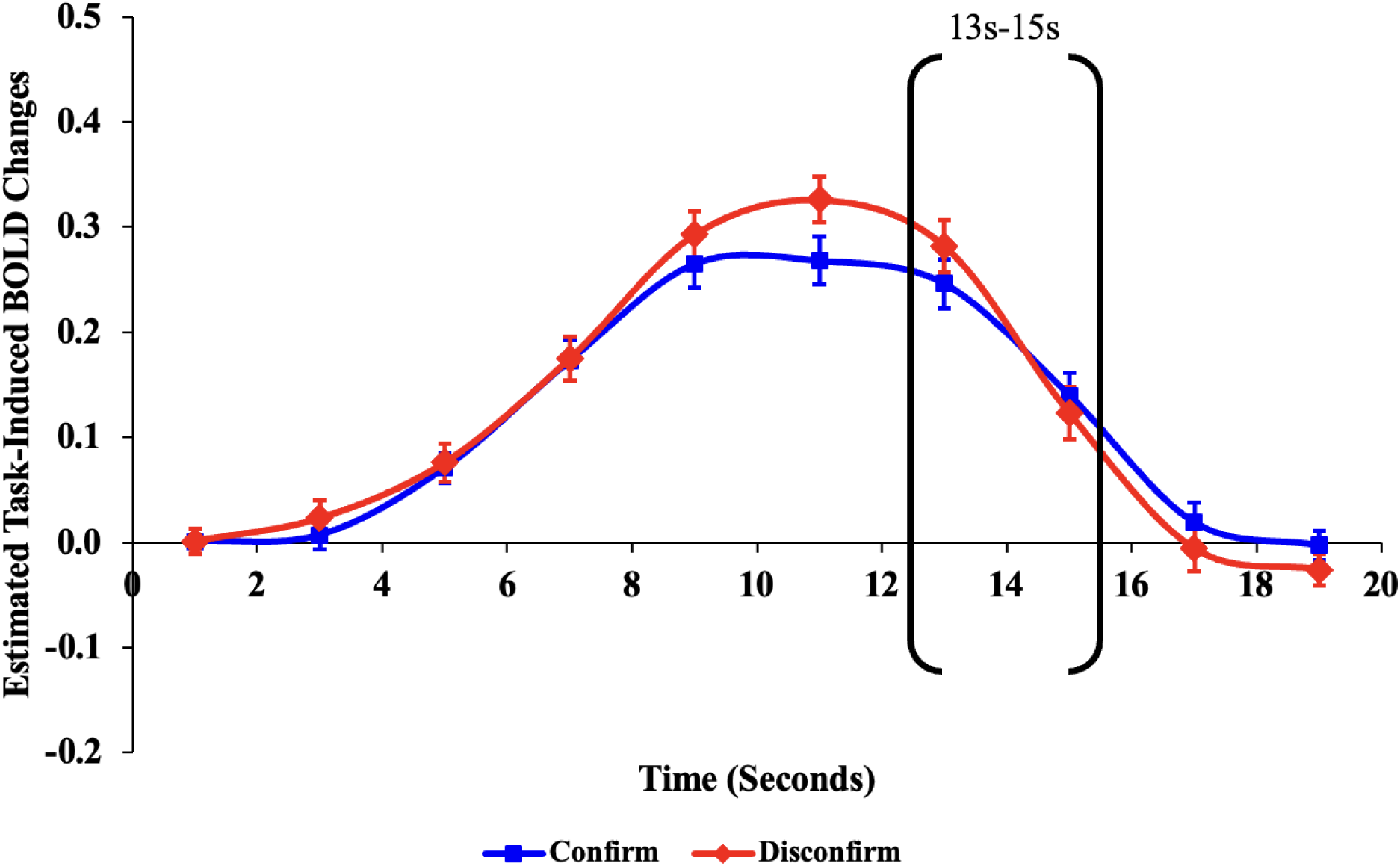
Estimated task-induced BOLD signal changes for PCBADE. Mean finite impulse response (FIR)-based predictor weights plotted as a function of post-stimulus time. Positive predictor weights plotted here reflect DMB deactivation, corresponding to the blue/green voxels in the associated anatomical patterns. Brackets indicate the task-induced BOLD changes dominating the Evidence Type × Time interaction, due to a higher peak for the disconfirm condition. The fMRI repetition time (TR) for this study was 2s.

#### 4.2.4. Raven’s Standard Progressive Matrices Task (RSPM)

A 3 (Difficulty - Easy, Medium, Hard) × 10 (Time) RM-ANOVA was conducted for the RSPM task, revealing a significant main effect of Time, *F*(9,495) = 74.87, *p* < .001, *η_p_^2^* = .58, and a significant Difficulty × Time interaction, *F*(18,990) = 2.35, *p* < .05, *η_p_^2^* = .04. This interaction was dominated by the contrast between 11.25s-13.75s, during which the Hard and Medium difficulty conditions demonstrated a steeper decline in DMB deactivation. This trend was a result of the Hard/Medium conditions showing a higher peak than the Easy condition at approximately 9s (see Figure 13), and a steeper decline to baseline.

**Figure 13.**
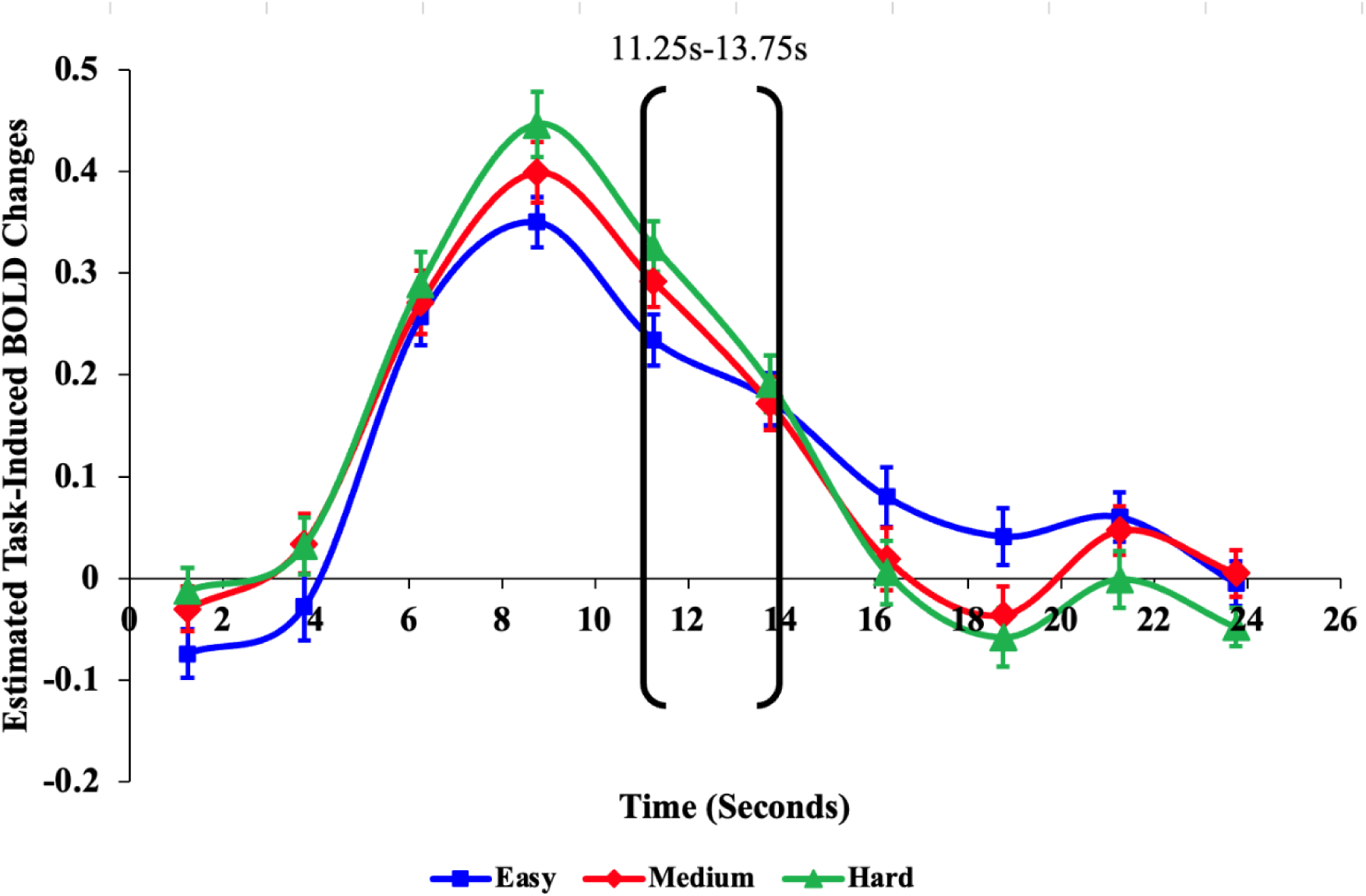
Estimated task-induced BOLD changes for the Raven’s Standard Progressive Matrices (RSPM). Mean finite impulse response (FIR)-based predictor weights plotted as a function of post-stimulus time. Positive predictor weights plotted here reflect DMB deactivation, corresponding to the blue/green voxels in the associated anatomical patterns. Brackets indicate the task-induced BOLD changes dominating the Time × Difficulty interaction due to a steeper decrease in deactivation in the hard and medium conditions compared to the easy condition. The fMRI repetition time (TR) for this study was 2.5s.

#### 4.2.5. Autobiographical Event Simulation (AES)

The 4 (Condition-Recall, Imagine-Past, Imagine-Future, Semantic Association) × 21 (Time) RM-ANOVA revealed a significant main effect of Time, *F*(20,320) = 4.26, *p* < .001, *η_p_^2^* = .21, and Condition, *F*(3,48) = 35.92, *p* < .001, *η_p_^2^* = .69, where the DMB showed activations in the autobiographical event simulation conditions (Figure 14, red, green, and yellow lines) and a strong deactivation in the Semantic Association condition (Figure 14, blue line). There was also a significant Condition × Time interaction, *F*(60,960) = 8.77, *p* < .001, *η_p_^2^* = .35 (Figure 14).

**Figure 14.**
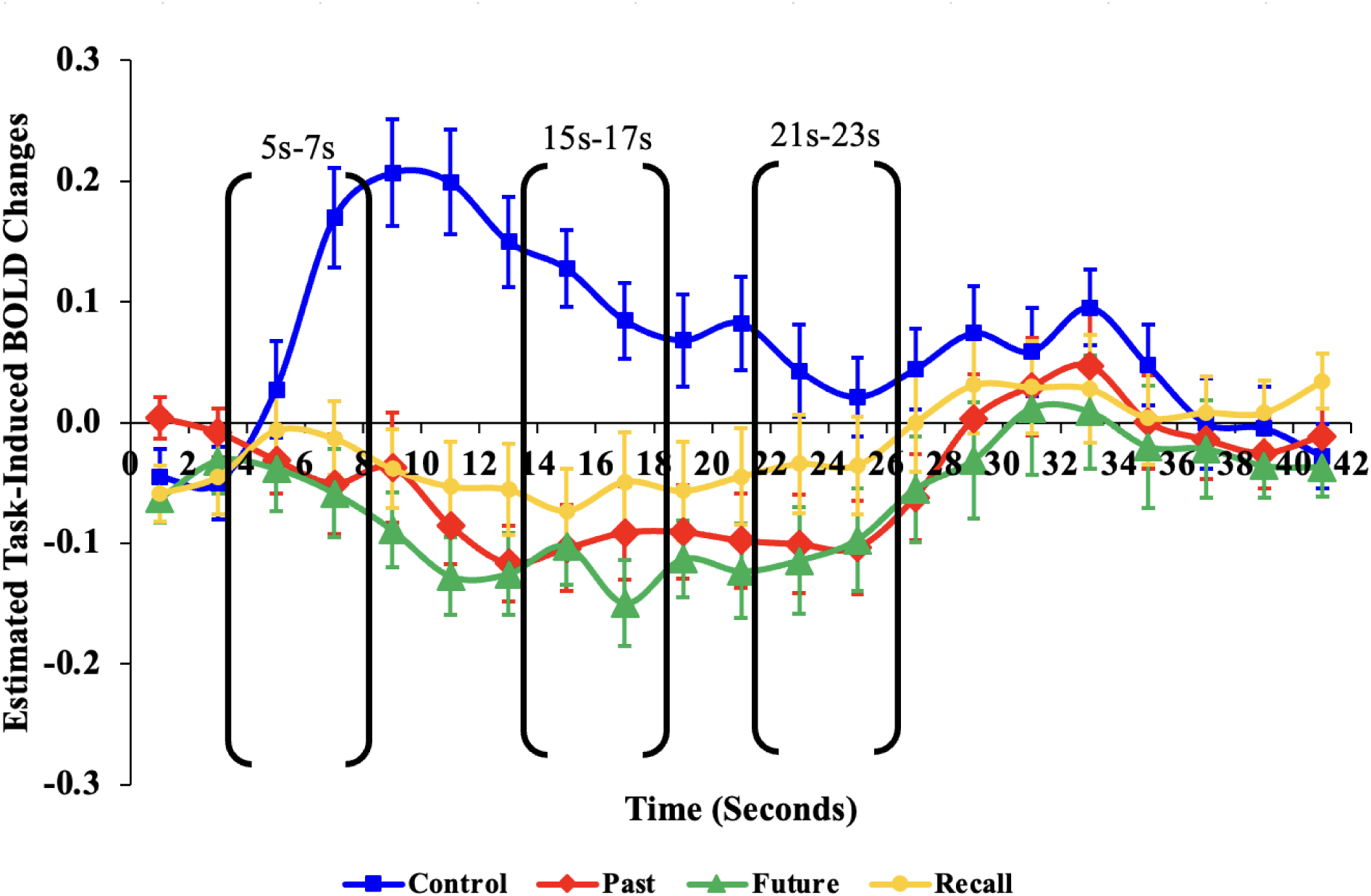
Estimated task-induced BOLD changes for the Autobiographical Event Simulation (AES). Mean finite impulse response (FIR)-based predictor weights plotted as a function of post-stimulus time. Positive predictor weights plotted here reflect DMB deactivation, corresponding to the blue/green voxels in the associated anatomical patterns. Whereas negative predictor weights reflect DMB activation. Brackets indicate the task-induced BOLD changes dominating the Time × Condition (Semantic Association, Imagine-Past/Recall) interaction due activations in the autobiographical event simulation conditions and deactivation in the semantic association condition. The fMRI repetition time (TR) for this study was 2s.

Using the difference contrasts, we found that DMB activity was greater in the Imagine-Future relative to the Imagine-Past condition, *F*(1, 16) = 6.42, *p* < .05, *η_p_^2^* = .29 (Figure 14, red *vs.* green lines). DMB activity in the Imagine-Past condition did not differ significantly from the Recall condition, *F*(1, 16) = 3.28, *p* = .09 (Figure 14, red vs. yellow lines), but the average activity of DMB in the Imagine-Past and Recall conditions was significantly higher than in the Semantic Association condition, *F*(1,16) = 41.42, *p* < .001, *η_p_^2^* = .72 (Figure 14, blue vs. yellow and red lines).

The Condition (Imagine-Future *vs.* Imagine-Past) × Time interaction was dominated by contrasts between 7s to 9s, *F*(1,16) = 7.02, *p* < .05, *η_p_ ^2^*= .31, and from 15s to 17s, *F*(1,16) = 4.86, *p* < .05, *η_p_^2^* = .23, where the DMB activity increased more steeply to peak activation in the Imagine-Future condition relative to the Imagine-Past condition (Figure 14, red vs. green lines).

The Condition (Semantic Association *vs*. Imagine-Past/Recall) × Time interaction was dominated by contrasts at three time points (Figure 14). Earlier in the trials, from 5s to 7s, *F(*1,16) = 51.00, *p* < .001, *η_p_ ^2^*= .76, the DMB activated in the autobiographical event simulation conditions but strongly deactivated in the Semantic Association condition (Figure 14, blue line vs. others). Later in the trials towards the end of the elaboration phase and beginning of the scale rating phase, from 15s to 17s, *F(*1,16) = 16.75, *p* < .001, *η_p_ ^2^* = .51, and from 21s to 23s, *F(*1,16) = 8.52, *p* < .05, *η_p_^2^* = .35, the activity of the DMB returned towards baseline in all conditions via deactivations and activation in the autobiographical event simulation and Semantic Association conditions, respectively (Figure 14, blue line vs. others).

#### 4.2.6. Mindfulness Meditation (MM)

Analyzing the Thought blocks, and excluding the Emotion condition due to a low frequency of selection, a 3 (Condition-Bodily, Narrative and Visual) × 18 (Time) RM-ANOVA revealed a significant main effect of Condition, *F*(2,30) = 5.82, *p* < .01, *η_p_^2^* = .28, whereby the DMB showed activations in the Narrative and Visual conditions (see Figure 15, red and green lines) and a deactivation in the Bodily condition (Figure 15, blue line), with no significant main effect of Time (*p* =.13). There was also a significant Condition × Time interaction, *F*(34,510) = 3.32, *p* < .001, *η_p_^2^* = .18 (Figure 15).

**Figure 15.**
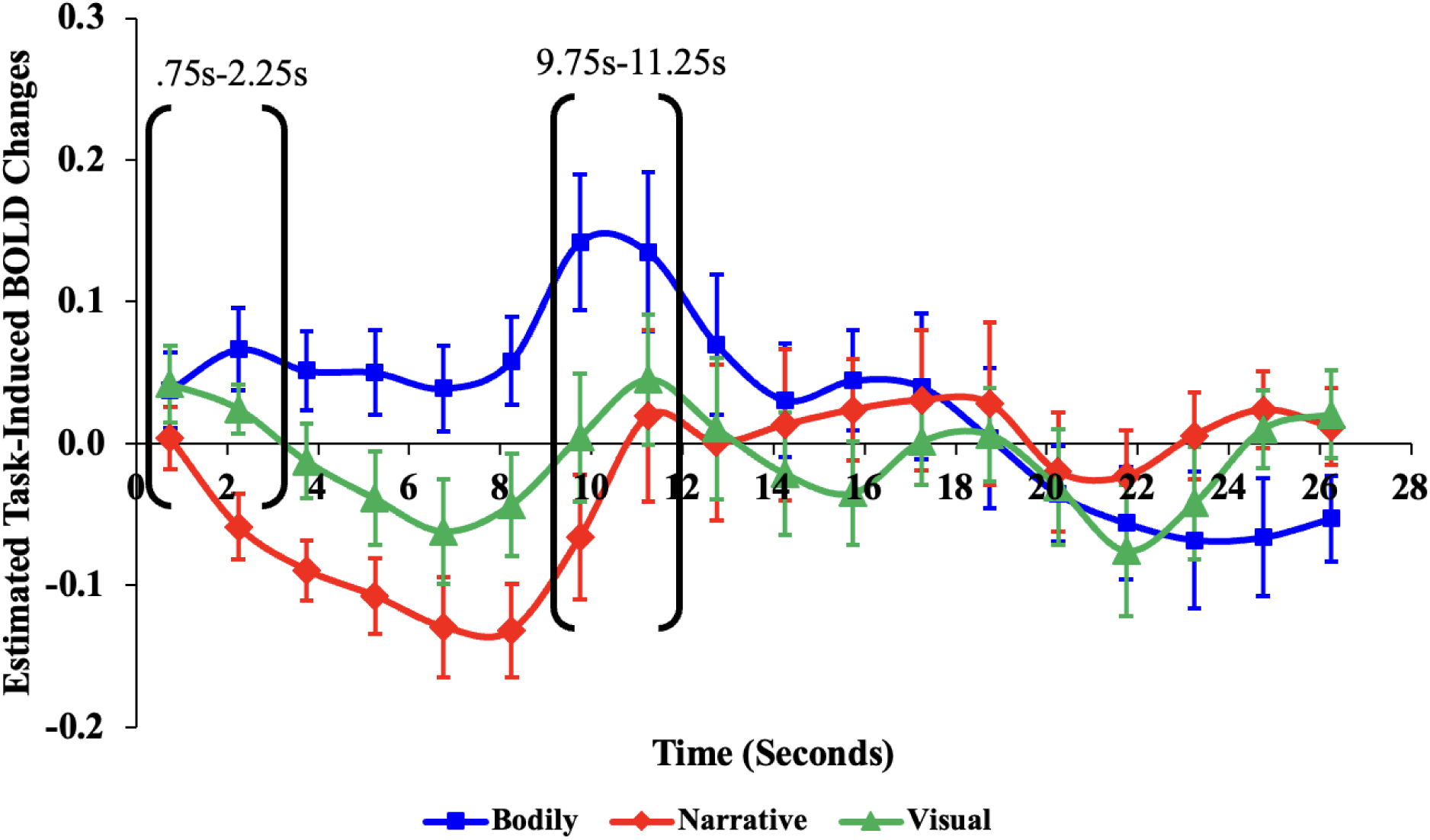
Estimated task-induced BOLD changes for the Mindfulness task. Mean finite impulse response (FIR)-based predictor weights plotted as a function of post-stimulus time. Positive predictor weights plotted here reflect DMB deactivation, corresponding to the blue/green voxels in the associated anatomical patterns. Whereas negative predictor weights reflect DMB activation. Brackets indicate the task-induced BOLD changes dominating the Time × Condition (Bodily vs. Narrative/Visual) interaction due activations in the Narrative and Visual conditions and deactivation in the Bodily condition. The fMRI repetition time (TR) for this study was 1.5s.

Using the difference contrasts, we found that DMB activity in the Narrative and Visual conditions did not differ (*p*=.23); however, the DMB activity in the Bodily condition significantly differed from its average activity in the Narrative and Visual conditions, *F*(1,15) = 7.41, *p* < .001, *η_p_^2^* = .33 (Figure 15, blue vs. others). The Condition (Narrative/Visual *vs.* Bodily) × Time interaction was dominated by two contrasts. Early in the trials, from.75s to 2.25s, *F*(1,15) = 9.16, *p* < .01, *η_p_^2^* = .38, the DMB activated in the Narrative and Visual conditions, but it deactivated in the Bodily condition (Figure 15, blue line vs. others). This led to the DMB proceeding to reach peak activations in the Narrative and Visual conditions but remaining below baseline in the Bodily condition. Subsequently, from 9.75s to 11.25s, *F*(1,15) = 10.42, *p* < .01, *η_p_ ^2^* = .41, the DMB showed a strong peak deactivation in the Bodily condition, but smaller peak deactivations in the Narrative and Visual conditions (Figure 15, blue line vs. others).

#### 4.2.7. Human Connectome Project - Social (HCP-Soc)

The 2 (Object Movement - Mental, Random) × 46 (Time) RM-ANOVA revealed a significant main effect of Time, *F* (45,22455) = 95.39, *p* < .001, *η_p_^2^* = .16, and Object Movement, *F* (1,499) = 29.16, *p* < .001, *η_p_^2^* = .06, where the DMB showed an activation in the Mental condition (see Figure 16, blue line) and a deactivation in the Random condition (Figure 16, red line). There was also a significant Object Movement × Time interaction, *F* (45,22455) = 40.54, *p* < .001, *η_p_^2^* = .08 (Figure 16).

**Figure 16.**
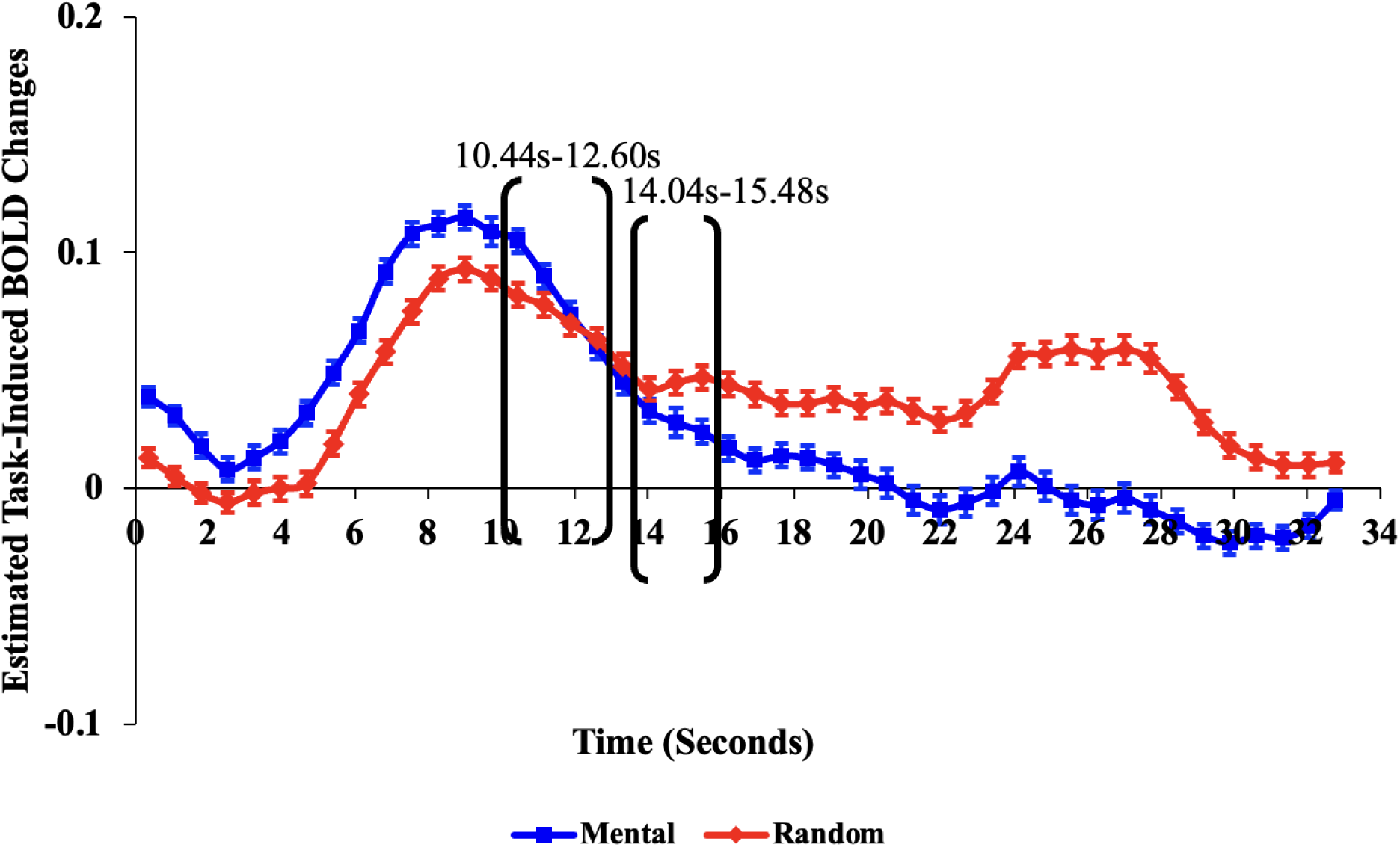
Estimated task-induced BOLD changes for Human Connectome Project - Social (HCP-Soc). Mean finite impulse response (FIR)-based predictor weights plotted as a function of post-stimulus time. Positive predictor weights plotted here reflect DMB deactivation, corresponding to the blue/green voxels in the associated anatomical patterns. Whereas negative predictor weights reflect DMB activation. Brackets indicate the task-induced BOLD changes dominating the Time × Object Movement interaction due to activation in the Mental condition and deactivation in the Random condition. The fMRI repetition time (TR) for this study was .72s.

The Object Movement (Mental, Random) × Time interaction was dominated by contrasts of adjacent time bins between 10.44s to 12.60s and 14.04s to 15.48s, due to initial deactivation of DMB early in the trial, followed by eventual activation, in the Mental condition relative to the Random condition (see Figure 16).

## 4. Discussion

In the context of task-based functional magnetic resonance imaging (fMRI), cognitive modes can be defined as task-general cognitive/sensory/motor processes which reliably elicit specific blood-oxygen-level-dependent (BOLD) signal pattern configurations. The function that defines the cognitive mode can be determined by study of changes in these BOLD signal patterns over a range of tasks and task conditions. Eight task studies with fMRI-CPCA-extracted components matching the spatial template of the DMB cognitive mode were analyzed and presented here. Their predictor weights were used to display the BOLD signal changes induced for each condition, and these BOLD signal changes were interpreted in the context of the DMB cognitive mode. Six out of eight tasks (SCAP, SA, PCBADE, RSPM, MM and HCP-Soc) were fully independent from the originally published DMB anatomical template image, and these independent studies replicated the patterns observed in the original anatomical template image. In addition, the interpretations of the independent studies supported the DMB account.

### 4.1. Working Memory with Varying Cognitive Load and Delay Length (WML46D04)

In the WML46D04 task, both increased working memory load and longer delay duration affected DMB deactivation. The increase in load led to the expected increased deactivation for 4-letter versus 6-letters conditions (see Figure 8, red > blue lines). When comparing the No Delay to a 4-second delay condition, the 4-second delay condition exhibited more sustained deactivation (see Figure 8, dotted lines extended relative to solid lines); however, unlike with the Load manipulation, the peak deactivation didn’t differ substantially between the delay conditions. This suggests that the DMB is not actually involved in coding the to-be-remembered material for maintenance, which can be contrasted with the mode referred to as maintaining attention to internal representations (MAIN) which does show higher activation for longer duration of delay for the WML46D04 task (Momeni et al., 2024; Sanford et al., 2020b), reflecting a cognitive mode that increases activity as more information is maintained.

### 4.2. Spatial Capacity (SCAP)

The deactivation of the spatial SCAP task replicated that of the verbal WML46D04 task, showing both increased deactivation for increased load (see Figure 9; 1 < 3 < 5 < 7 dots), and longer delay conditions exhibited more sustained deactivation (see Figure 9; green extended relative to red, and red extended relative to blue), but unlike the Load manipulation, the peak deactivation didn’t differ substantially between the delay conditions. This confirms that the DMB is not actually involved in coding the to-be-remembered material for maintenance, which can be contrasted with the MAIN mode, which does show higher activation for longer duration of delay for the SCAP task (Momeni et al., 2024).

### 4.3. Semantic Association (SA)

In the SA task, both conditions showed deactivation of the DMB, with the Distant condition exhibiting greater DMB deactivation than the Close condition (Figure 11, red line *>* blue line). In the Distant condition, selecting a word distantly related to the prompt word requires more attentional resources, resulting in stronger DMB deactivation, a replication of the effect of load dependence.

### 4.4. Picture Completion Bias Against Disconfirmatory Evidence (PCBADE)

In the PCBADE task, both conditions showed deactivation of the DMB, with the Disconfirm condition exhibiting greater DMB deactivation than the Confirm condition (Figure 12, red line *>* blue line), as the Disconfirm condition required more attentional resources, resulting in stronger DMB deactivation, a replication of the effect of load dependence.

### 4.5. Raven’s Standard Progressive Matrices (RSPM)

In the RSPM task, more difficult conditions elicited greater DMB deactivation.

Specifically, the Hard difficulty condition exhibited the highest peak of deactivation, followed by the Medium condition, and the Easy condition (Figure 13 green line *>* red line *>* blue line), a replication of the effect of load dependence.

### 4.6. Autobiographical Event Simulation (AES)

In the AES task, DMB showed a strong deactivation in the semantic association condition (Figure 14, blue line *vs.* others), replicating the results of the SA task presented above. In sharp contrast, the DMB showed activations in all three autobiographical event simulation conditions (Figure 14, green, red and yellow line *vs.* blue), an effect noted in previous studies (Addis et al., 2009; Benoit & Schacter, 2015). This clear contrast demonstrates that even within one task, instructions/stimuli can have a strong effect on whether or not the DMB activates or deactivates. The cognitive difference between semantic association *versus* all three autobiographical event simulation conditions, but also a cognitive process being common to imaging the future, past and recall of past, could be mental projection into self-relevant social narratives. However, it also must be noted that DMB was not the only cognitive mode to be activated during autobiographical event simulation, as the MAIN and multiple demand (MD) modes also activated, as noted elsewhere (Momeni et al., 2025; Momeni et al., 2024; Wang et al., 2024).

However, it is important to keep in mind that under most task condition, with autobiographical event simulation being a notable exception, DMB deactivates while MAIN or MD activate (Momeni et al., 2024; Wang et al., 2024). This reciprocal relationship between DMB on one hand, and MAIN/MD on the other, is observed in the semantic association condition reported here with the AES task, but not the three autobiographical event simulation conditions in the AES task.

### 4.7. Mindfulness Meditation (MM)

In the Mindfulness task, DMB showed deactivation in the Bodily condition, but activations in the Visual and Narrative conditions (Figure 15, blue line peak > 0 at ∼11s, *vs.*green/red lines peak < 0 at ∼7s). The DMB network has been previously shown to activate during generation of spontaneous thoughts (Christoff et al., 2016). Thus, the activations of the DMB in the Visual and Narrative conditions align with previous research. The activation of the DMB in the Bodily condition, unlike the other conditions, would be expected based on the assertion that DMB deactivates during attentional focus on external sensory experiences, as opposed to engaging in mental projection into self-relevant social narratives, such as in the Visual and Narrative conditions, as would be expected for attention to features of the external environment (Buckner, 2013).

### 4.8. Human Connectome Project – Social (HCP-Soc)

In the HCP-Soc task, early in the trials, the DMB showed deactivations in both the Mental and Random conditions, likely due to participants directing their attention externally to the movie stimuli when they were first presented. However, as the trial progressed, the DMB deactivation diverged between the Random and Mental conditions, with the Mental condition being pushed towards activation (i.e., < 0), as participants judged that the movement of objects depicted a social interaction (Figure 16, blue line trajectory < red line trajectory after 14s).

Previous research has shown that the core default mode activates during fMRI tasks that engage theory of mind (i.e., taking another’s perspective; Buckner et al., 2008a; DuPre et al., 2016; Tamir et al., 2016). The results from the HCP-Soc task align with these findings as in the Mental condition, participants engaged in theory of mind to evaluate the social interaction among the movie stimuli, not elicited in the Random conditions. Finding a cognitive process that accords with the observations above, and for activations in AES and MM tasks, we can converge on engaging in mental projection into self-relevant social narratives. Although in this case the theory of mind task is not specifically referencing events in the autobiographical trajectory of the participant themselves, as was the case with AES and MM, DMB activation may extend to the types of social interactions that the participant has experienced themselves in the past, even if that experience is only observed in others’ actions currently.

### 4.9. Comparison to Other Networks and Cognitive Modes

It is important to note that all of the DMB results presented here were originally extracted in the context of two to seven other modes involved in the same task. Although anatomical comparisons of DMB to the other known cognitive modes are available for observation (Percival et al., 2020), it is the temporal comparison between cognitive modes which provides the evidence for the cognitive modes causing the set of patterns to emerge. These task-induced BOLD-signal change comparisons made across cognitive modes, and also within tasks, provide strong evidence for the functional mode interpretations. These are not addressed in this manuscript, but form the foundation of the cognitive mode approach to brain imaging (Fouladirad et al., 2022; Gill et al., 2022; Kusi et al., 2022; Lavigne, Menon, Moritz, et al., 2020; Lavigne, Menon, & Woodward, 2020; Momeni et al., 2025; Sanford et al., 2020a, 2020c; Sanford & Woodward, 2021a, 2021b; Wong et al., 2020; Zurrin et al., 2024).

Alongside DMB, there exists another task-negative cognitive mode termed Default Mode A (DMA; Jian et al., 2024). Here we provide a comparison between these two “default” cognitive modes. This comparison is crucial because DMA and DMB are often considered the same due to their task-negative aspects and similar neuroanatomy. However, a head-to-head comparison shows their unique cognitive functions and anatomical patterns. To clarify their distinction, we compare the anatomical regions involved in DMA and DMB using prototype slices comparing each mode (see Table 11 and Table 12).

**Table 11.**
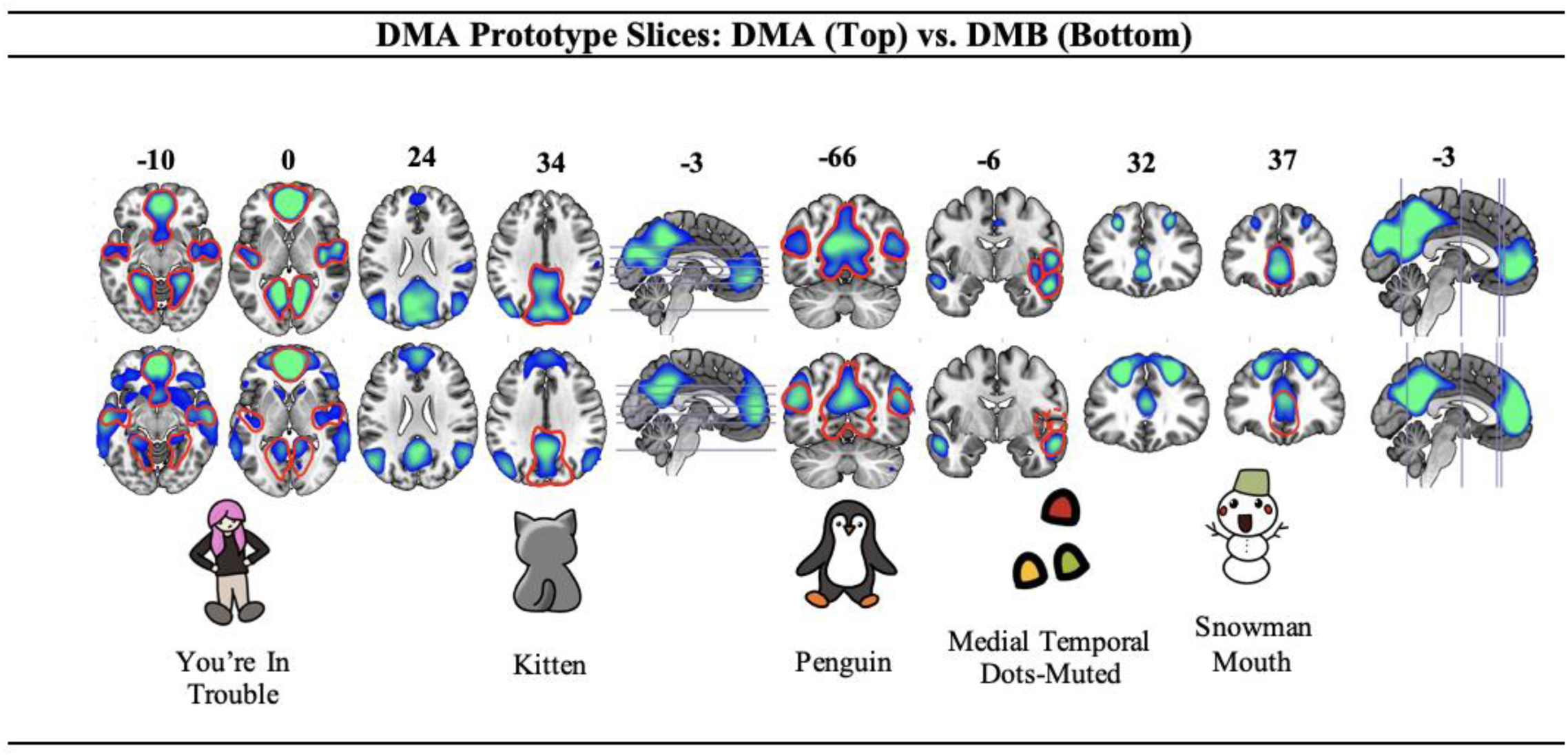
Prototype slices for the Default Mode A (DMA), comparing DMA (top) with the Default Mode B (DMB) (bottom).

**Table 12.**
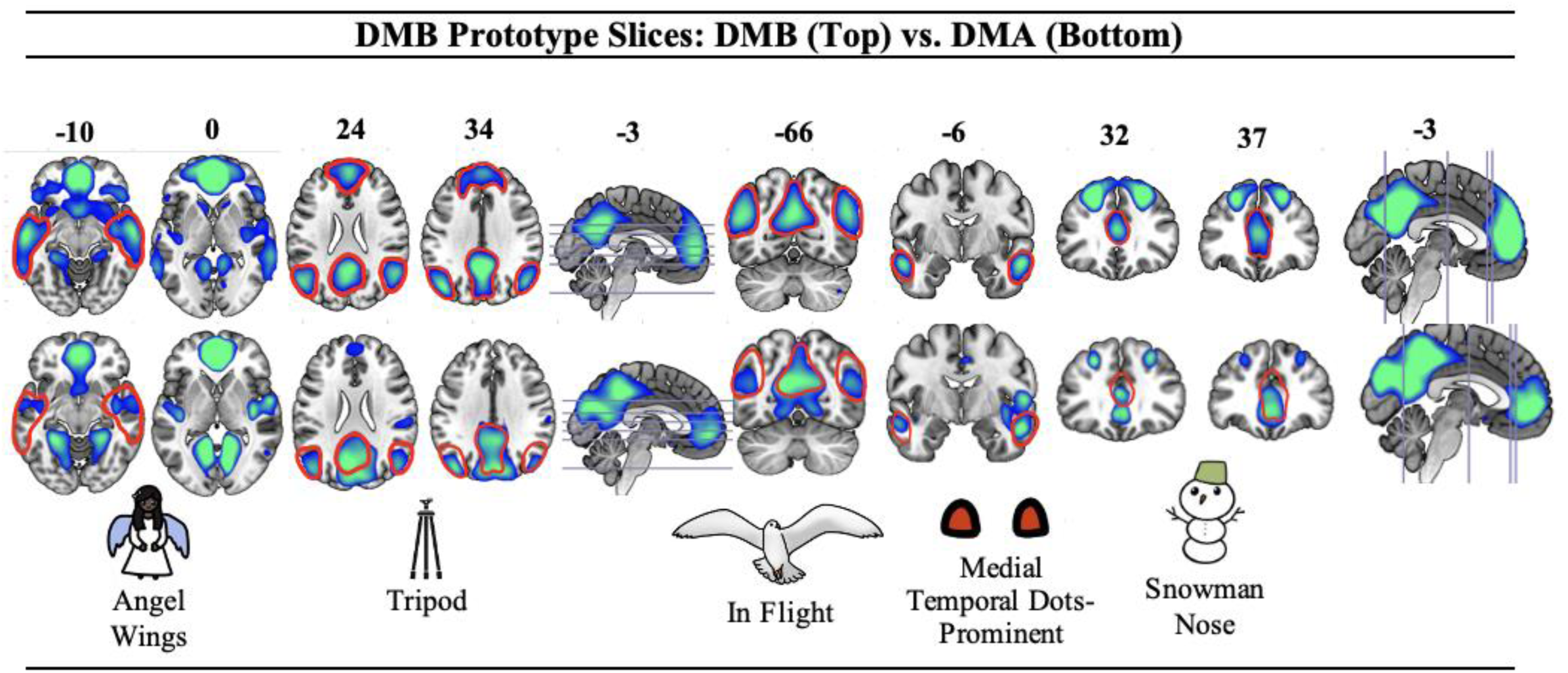
Prototype slices for the Default Mode B (DMB), comparing the DMB (top) with the DMA (bottom).

For example, in Table 11, axial slice 34, the DMA “Kitten” pattern (top row) shows deactivation in the posterior cingulate cortex and the precuneus, similar to DMB. However, DMB (bottom row) lacks the “haunches” of the Kitten, shown in the cuneus region. Comparing to Table 12, slice 34, the middle leg of the DMB “Tripod” pattern (top row) is extended to the “Kitten” haunches on DMA (bottom row). The headpiece of the tripod, present in the superior frontal gyrus for DMB (Table 12, top row slice 34), is absent in DMA (Table 12, bottom row slice 34). In addition, in axial slices -10 and 0, both DMA and DMB show prominent deactivation forming the “You’re in Trouble” and “Angel Wing” patterns, respectively.

However, the bilateral fusiform deactivation forming the “feet” of the DMA “You’re in Trouble” pattern (Table 11, axial slices -10 and 0 top row) is less pronounced in DMB (Table 11, axial slices -10 and 0 bottom row), and the wings of the DMB “Angel Wing” pattern in the Bilateral basal ganglia (Table 12, axial slice -10 top row) are not distinct in DMA (Table 12, axial slice - 10 bottom row).

In coronal slice -66, the DMA “Penguin” pattern’s legs in the lingual gyri (Table 11 top row, coronal -66) are not present in DMB (Table 11, bottom row, coronal -66). In coronal slice 37, the “Snowman Mouth” pattern in DMA (Table 11, top row, coronal 37) overlaps with the “Snowman Nose” pattern in DMB (Table 12, top row, coronal 32 and 37), but DMA’s “Snowman Mouth” deactivation is more inferior than the “Snowman Nose” (Table 11, top *vs.* bottom row, coronal 37; Table 12, top *vs.* bottom row, coronal 37). The DMB pattern “Medial

Temporal Dots-Prominent” share similar regions of deactivation between DMA and DMB, except the posterior part of the left Middle Temporal Gyrus is not present in the DMA deactivation. (see Table 11 and Table 12, top *vs.* bottom row, coronal -6).

Moving from anatomical to temporal activation patterns, together with the well-established literature on DMB (Andrews-Hanna et al., 2010; Buckner et al., 2008b; Raichle et al., 2001), the observed examples of both activation and load-dependent deactivation in this manuscript help specify DMB’s role in engaging in mental projection into self-relevant social narratives, and load dependent deactivation when attending to behaviourally relevant features of the external environment (Buckner, 2013). In contrast, default mode A (DMA; Jian et al., 2024) is a relatively novel cognitive mode with less understood functional and temporal dynamics. It has been observed the DMA is consistently deactivated reciprocally to the language mode (LAN; Zeng et al., 2024); however, there is no record of activation to aid with interpretation as is the case with DMB.

Comparisons of DMB to the 7-network Yeo resting-state templates have been published in the supplementary material of several recently published papers; namely, for the SCAP task (Sanford, 2019, Figure 4.10), PCBADE task (Lavigne, Menon, Moritz, et al., 2020, Supplementary Table S2), RSPM task (Zurrin et al., 2024, Supplementary Table S1) and AES task (Momeni et al., 2025, Appendix Table A7). In Appendix Figure A1 we provide the Yeo/Buckner/Choi networks (Buckner et al., 2011; Choi et al., 2012; Yeo et al., 2011) on which each of the DMB specific pattern peaks are located. Of these 22 peaks, 19 were located on the default-mode network, one on the limbic network, and two on the visual network.

Correspondingly, in Appendix Figure A1 we see that the primary overlap was with the “default-mode” network (Kong et al., 2024).

It is also instructive to compare the function of DMB to that of MAIN (Momeni et al., 2025; Momeni et al., 2024; Sanford et al., 2020b), since both may be thought to be involved in “internal thought”. As has been noted above, DMB deactivates during the type of “internal thought” involved in coding to-be-remembered material for maintenance during working memory or semantic association, but activates during the type of “internal thought” involved in engaging in mental projection into self-relevant social narratives. This comparison highlights the importance of specific functional definitions beyond general terms such as “internal thought”.

### 5.12. Limitations

The major limitation to the work was that two (WML46D04-TG and AES) of the eight tasks reported here had overlapped with the originally available anatomical template image (Percival et al., 2020). However, the other six were fully independent (SCAP, SA, PCBADE, RSPM, MM and HCP-Soc), and the independent studies replicated very well the key anatomical patterns observed in the original anatomical template image (see Table 2). Importantly, the interpretations of the task-induced BOLD-signal changes in the fully independent studies were highly reliable in their support of the DMB cognitive mode account, both anatomically and functionally.

It is important to note that these eight tasks were not only very different with respect to timing, stimuli, and objectives (e.g., clinical vs. non-clinical studies), they were collected in five different cities. WML46D04-TG, PCBADE, SA, and MM were collected in Vancouver, Canada (Philips 3T MRI scanner; Woodward PI). SCAP was collected in Los Angeles, USA (two Siemens Trio 2T MRI scanners; Poldrack PI). RSPM was collected in Calgary, Canada (General Electric Discovery MR750 3T MRI scanner; Goghari PI). AES was collected in Boston, USA (Siemens Sonata 1.5T MRI scanner; Addis PI). HCP-Soc was collected in Washington, USA (Siemens 3T scanner; Barch PI). Thus, the reliability of the anatomical patterns and the functional interpretations were subjected to the very stringent test of replicating over not only tasks and samples but also over data collection sites, scanners and investigators.

In future work, it will be necessary to carry out more completely independent analyses with new data and larger samples. Observation of naturally occurring configurations allow the determination of which prototypical patterns replicate or require adjustment/expansion. Also, as fMRI methods advance, with faster repetition times (TRs), and more spatial precision (e.g., layer fMRI), we may be able to expand the library of cognitive modes.

A perceived limitation of this work may be that this approach requires a degree of reverse inference. Reverse inference can be defined as the assignment of a cognitive function to an anatomical pattern. This term elicits caution for many cognitive neuroscientists because it requires something close to one-to-one mapping between the anatomical pattern on one hand, and the proposed cognitive operation on the other (Poldrack, 2006). This is widely considered to be virtually impossible, because both task-based brain activations (neurons) and the operations that are the building blocks of the cognitive architecture (bequeathed from cognitive psychology), are arguably infinite, and no classification method could enable both to be arranged into a finite set of units that could be directly linked. For example, neural activity associated with tasks has been described as “just a transient and task-specific association of brain regions” (Power et al., 2011).

However, task-based BOLD patterns bypass much of this complexity. fMRI does not directly measure any property of neural firing, but instead measures fluctuations in BOLD signal, and the correlated (over time) BOLD signal appears to reliably arrange into a relatively small set of patterns. Here we present anatomical configurations/patterns that reliably relate to the task-general cognitive mode of DMB. The “objective discovery” approach for brain function calls for an objective fMRI-specific vocabulary to replace/refine a vocabulary inherited from philosophy and psychology (Buzsaki, 2020). Classification of anatomical patterns, as well as interpretation of the temporal profiles of BOLD signal changes, followed by assignment of cognitive functions that define the mode, moves towards a nearly one-to-one mapping between BOLD network patterns and fMRI-specific task general cognitive processes. This method provides a potentially tractable solution to reverse-inference hesitancy inherent to task-based fMRI, and could bring more of the full potential of fMRI for clinical assessment of cognitive function within reach.

## 5. Conclusion

The correlated, time-dependent, task-induced BOLD signal changes consistently arrange into a relatively small set of anatomical patterns with an associated cognitive mode. DMB is one such mode, and it has been shown to be anatomically reliable. Based on observation of task-induced BOLD signal changes across a wide range of tasks, it can be observed that deactivations in DMB are sensitive to cognitive load during externally directed tasks, which require attention to specific behaviourally relevant features of the external environment. This was observed during WML46D04, SCAP, SA, PCBADE, and RSPM tasks. In contrast, MM, HCP-Soc, and AES showed activations in DMB. All could be interpreted as involving a cognitive process for engaging in mental projection into self-relevant social narratives. This includes theory of mind (HCP-Soc), imagining/remembering autobiographical memories intentionally (AES), and through the types of self-relevant social narratives that occur during mind wandering (MM).

Thus, deactivations in DMB were sensitive to cognitive load during attention to specific features of the external environment, and activations in DMB involved a cognitive process for engaging in mental projection into self-relevant social narratives. Future research may explore DMB activation over a wider range of tasks.

## Acknowledgements

We thank John Shahki who worked on aspects of this chapter. We thank Drs. Eva Feredoes, Daniel Margulies and Sepideh Sadaghiani for discussions which shaped the cognitive mode perspective. We also express our gratitude and acknowledge the financial and personal resources committed by our valued colleagues who procured fMRI data and/or contributed to the published work on which this manuscript was based, as listed in citations of published papers throughout the text.

## Supplementary Materials

**Supplementary Table S1.**
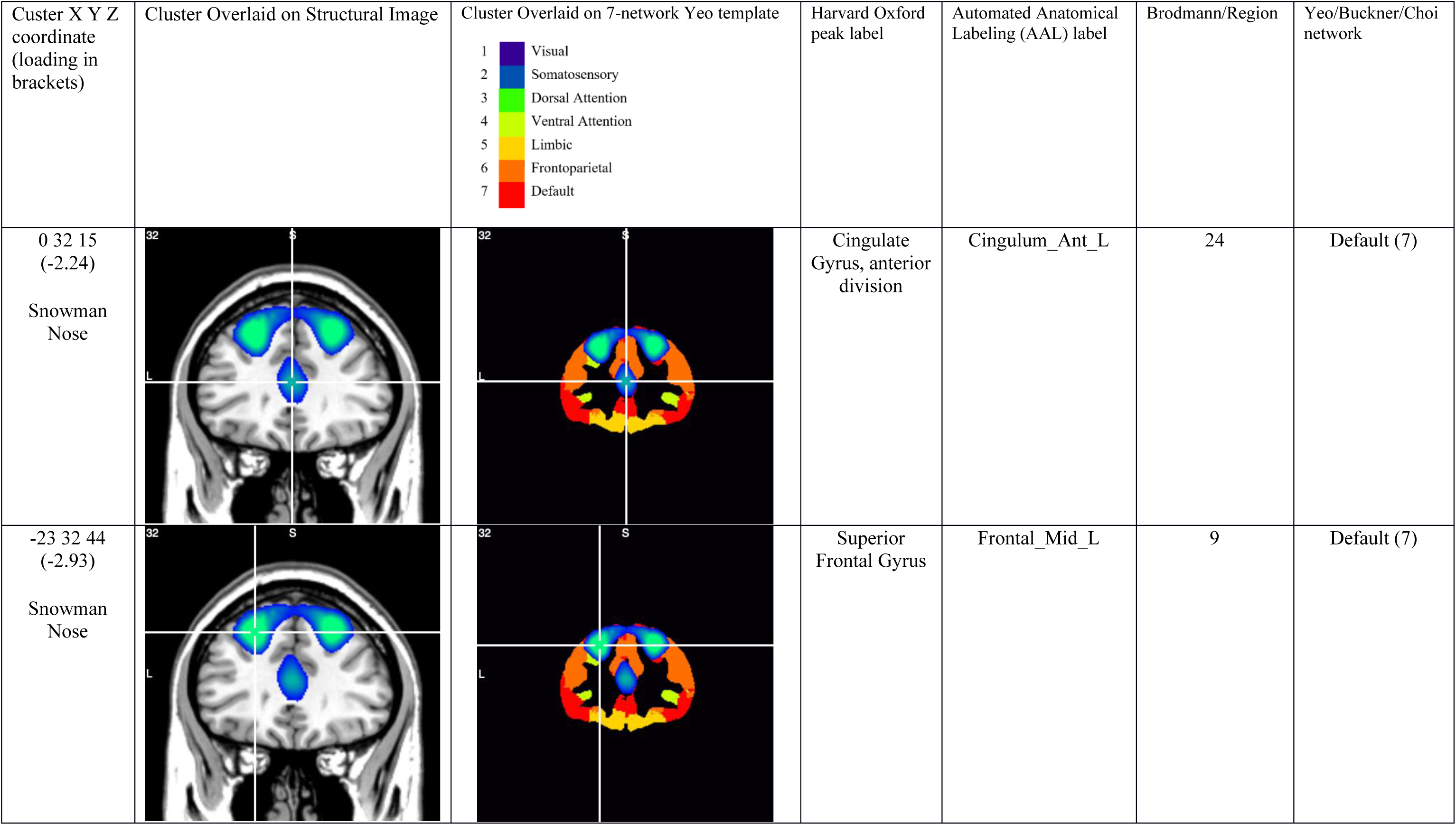
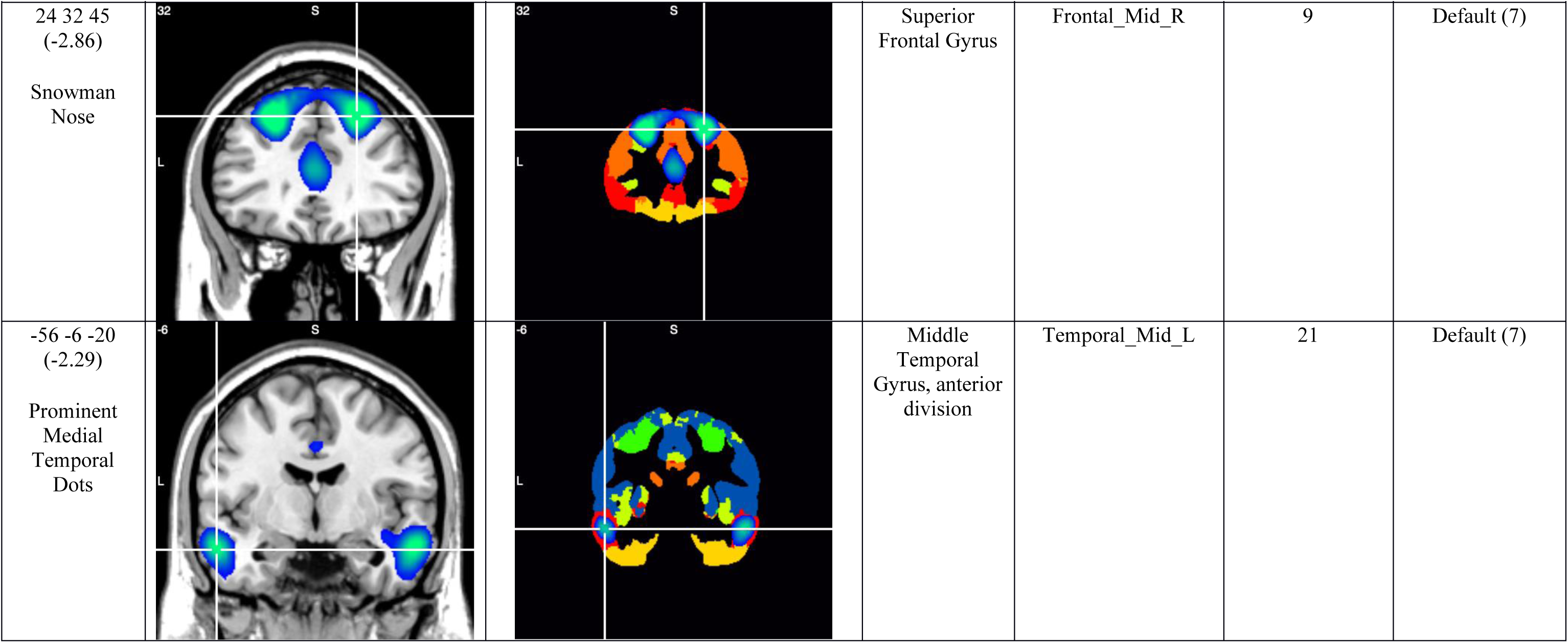
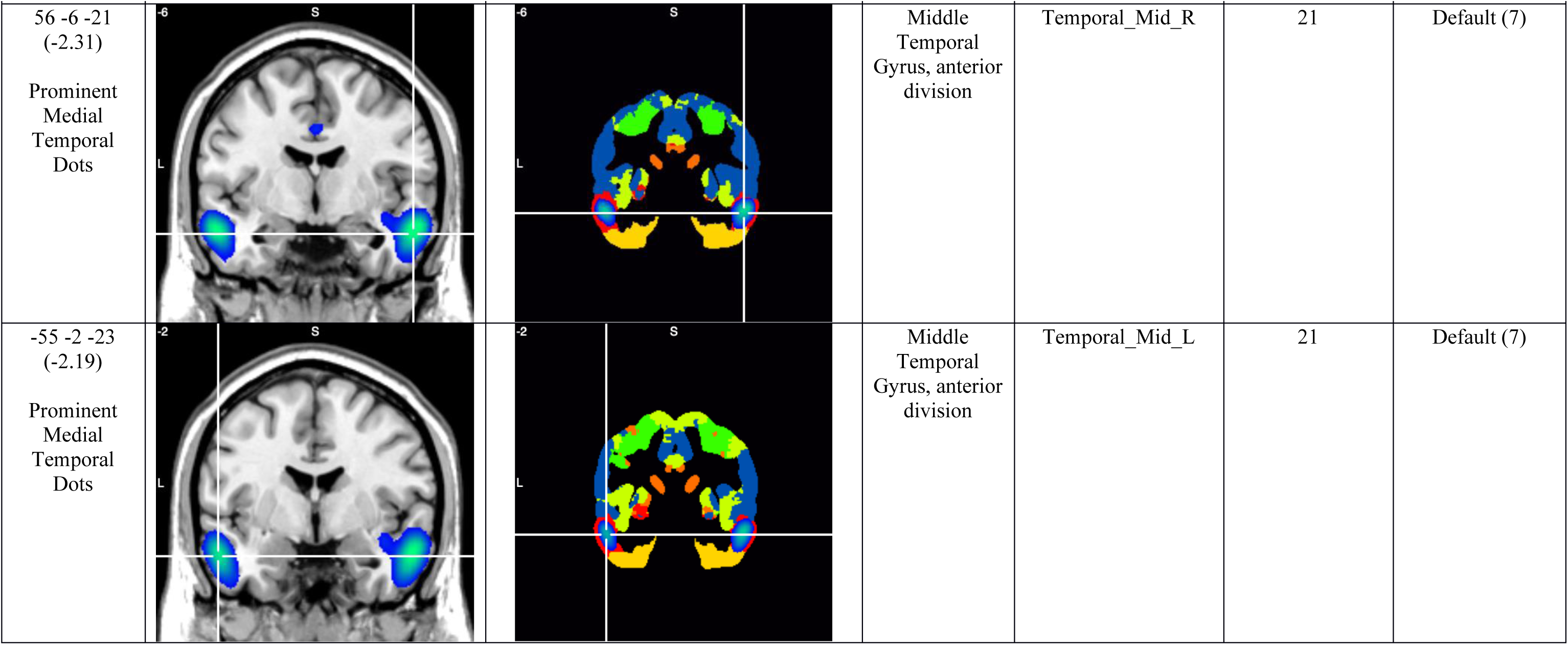
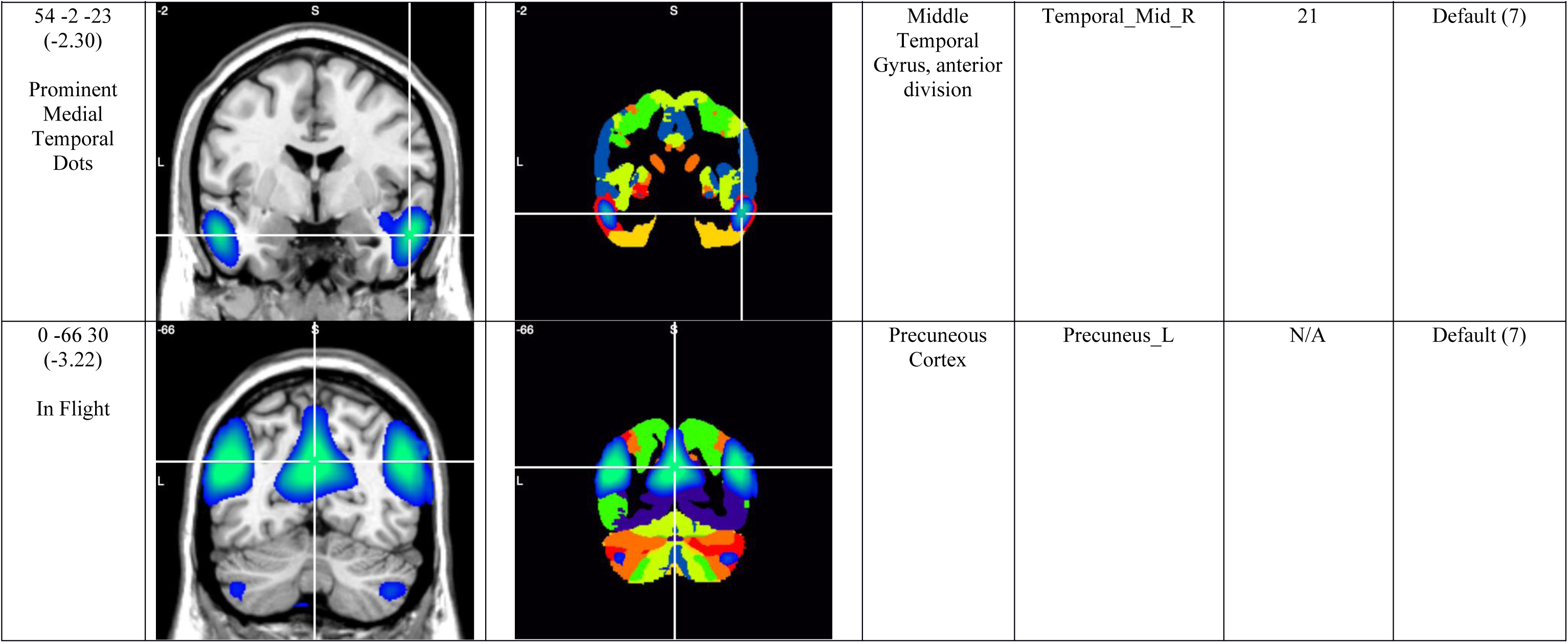
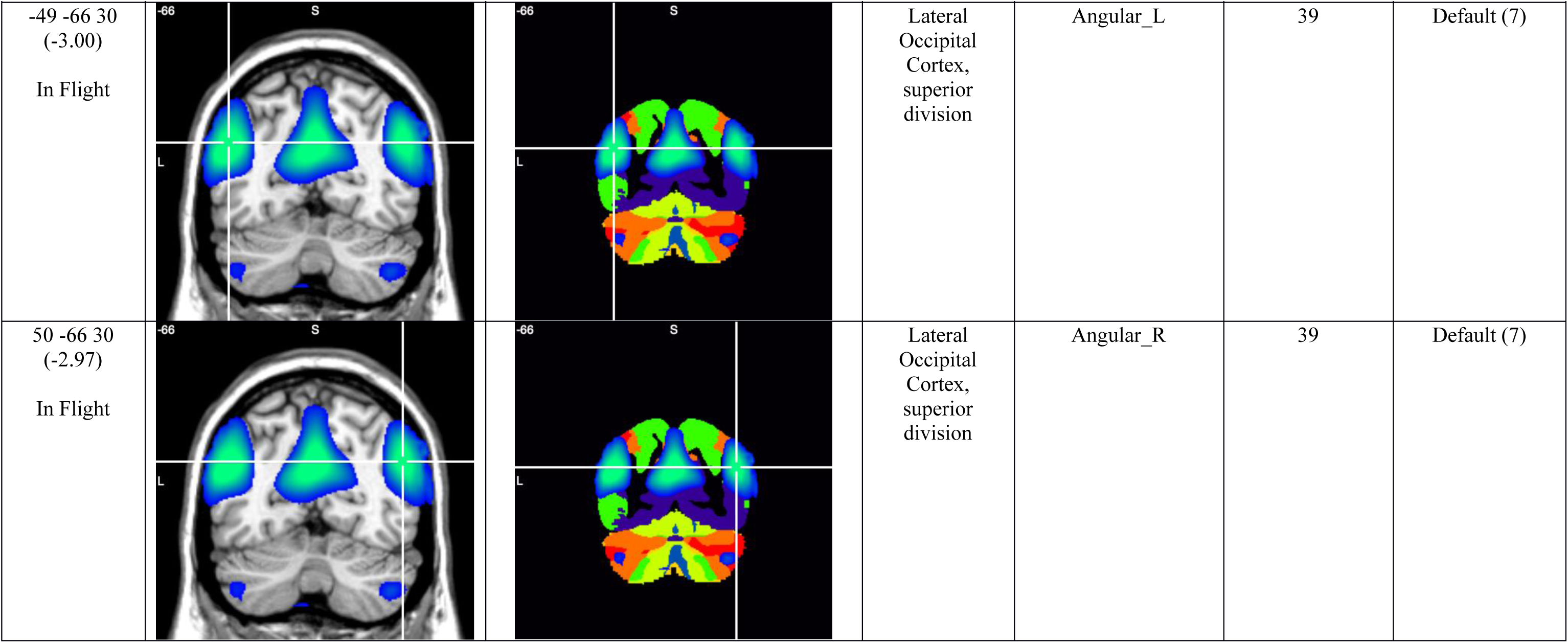
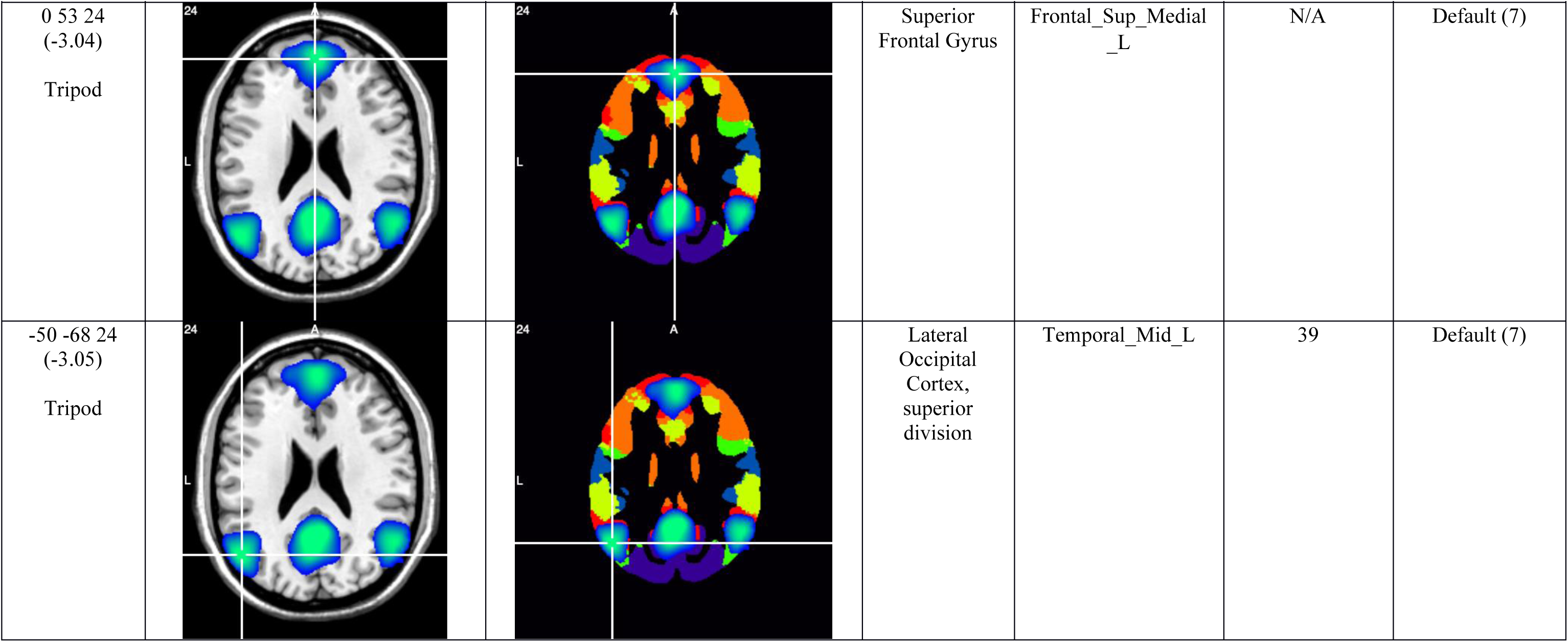
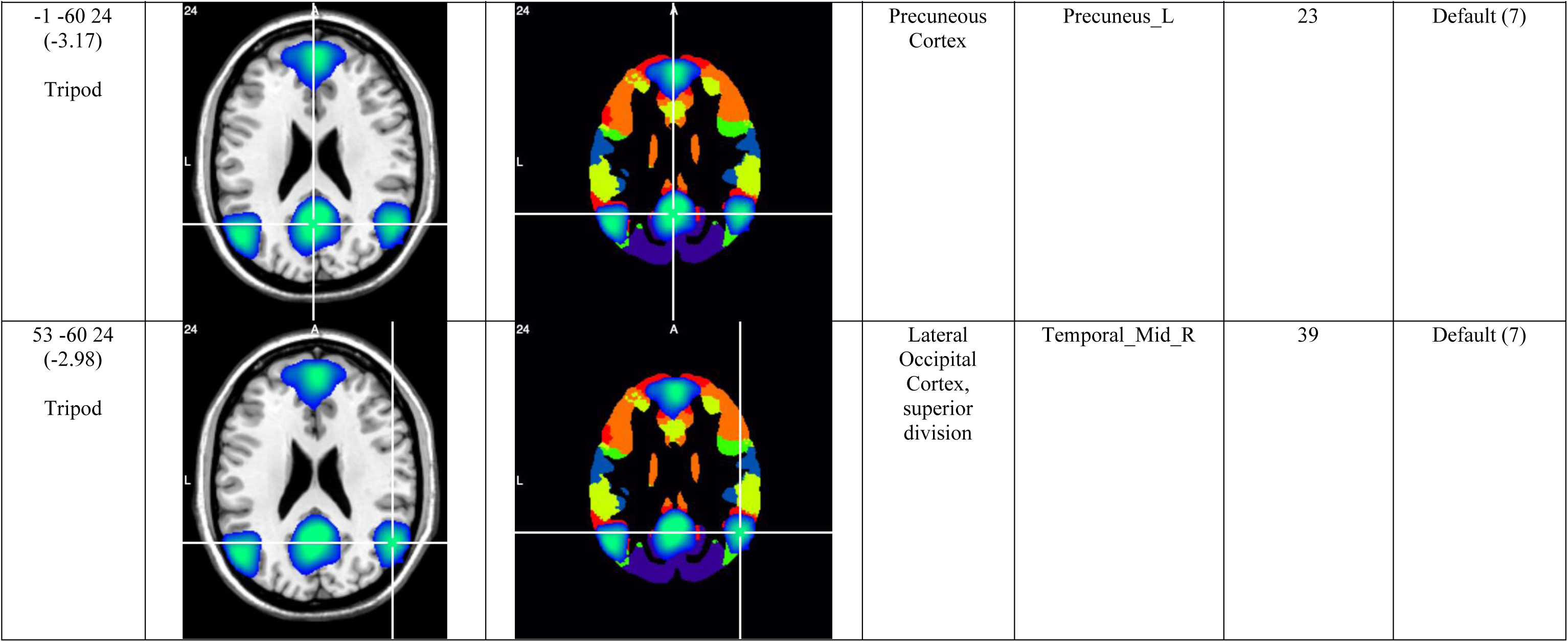
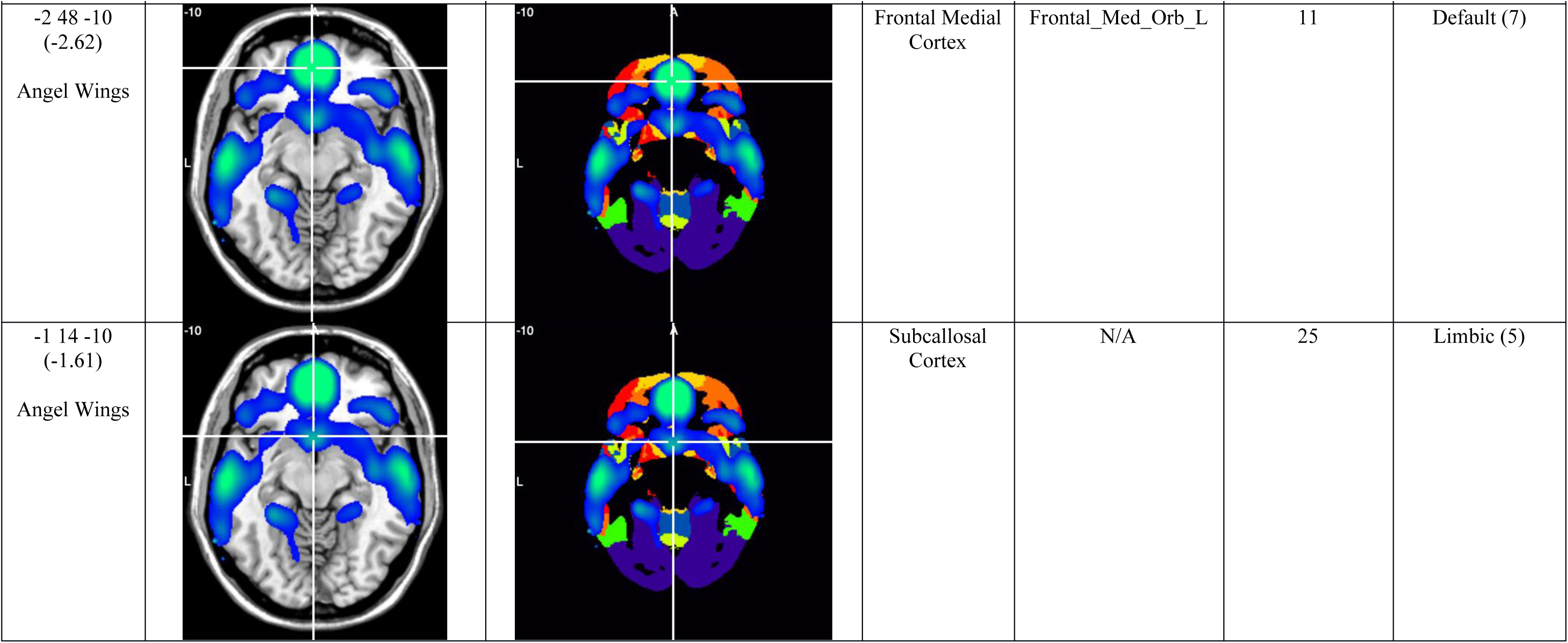
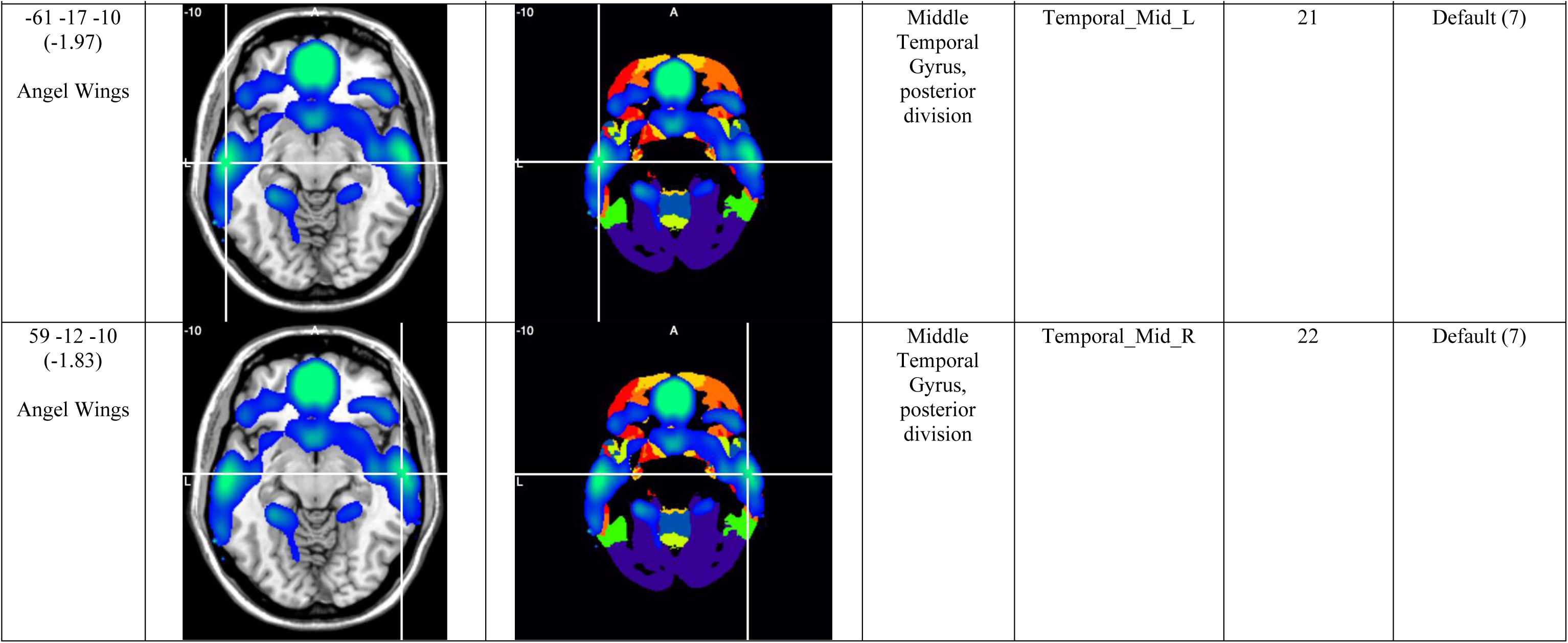
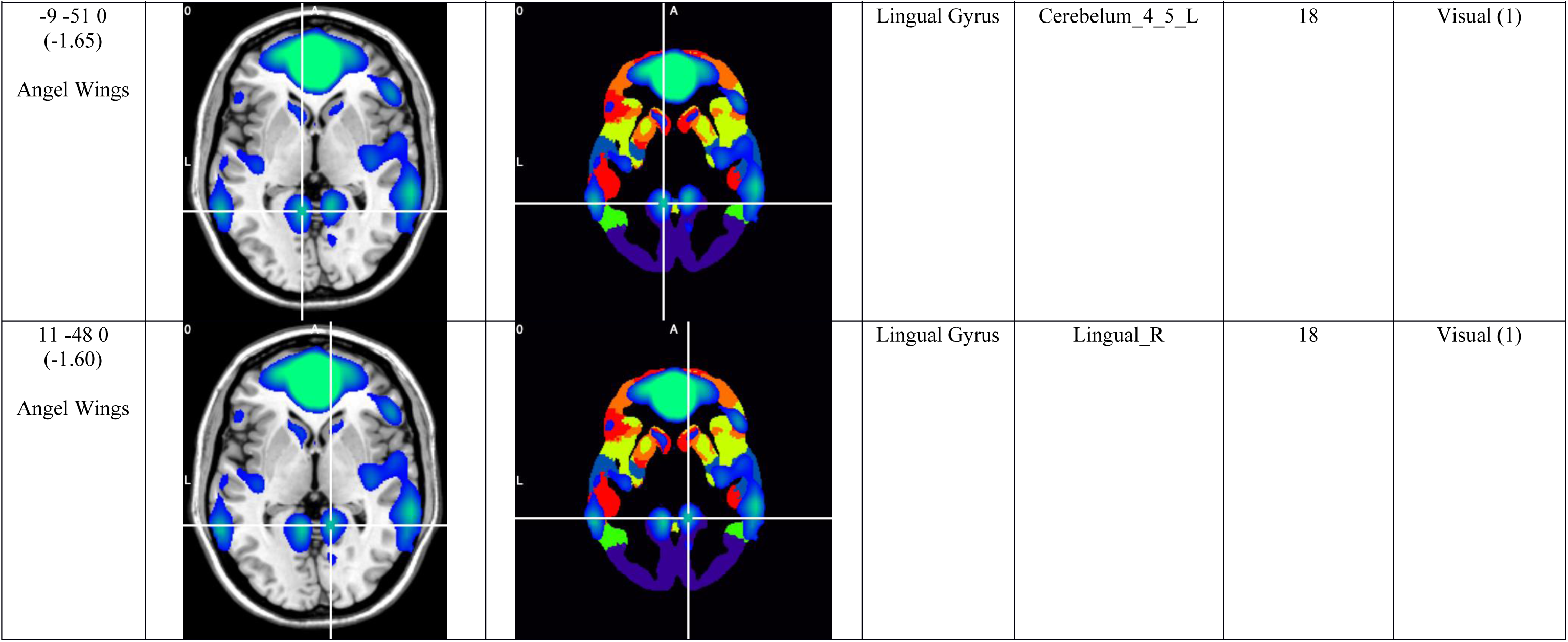
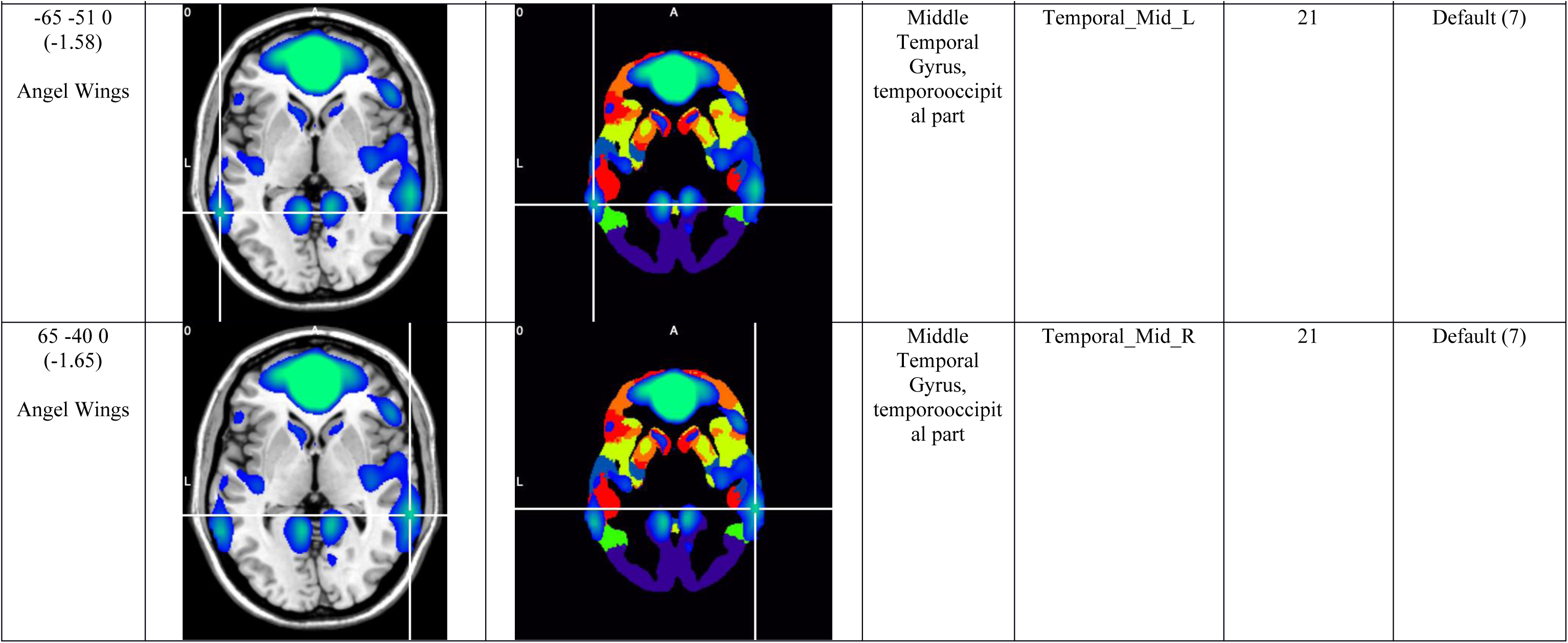
Default Mode B Mean Image.

**Appendix Figure A1.**
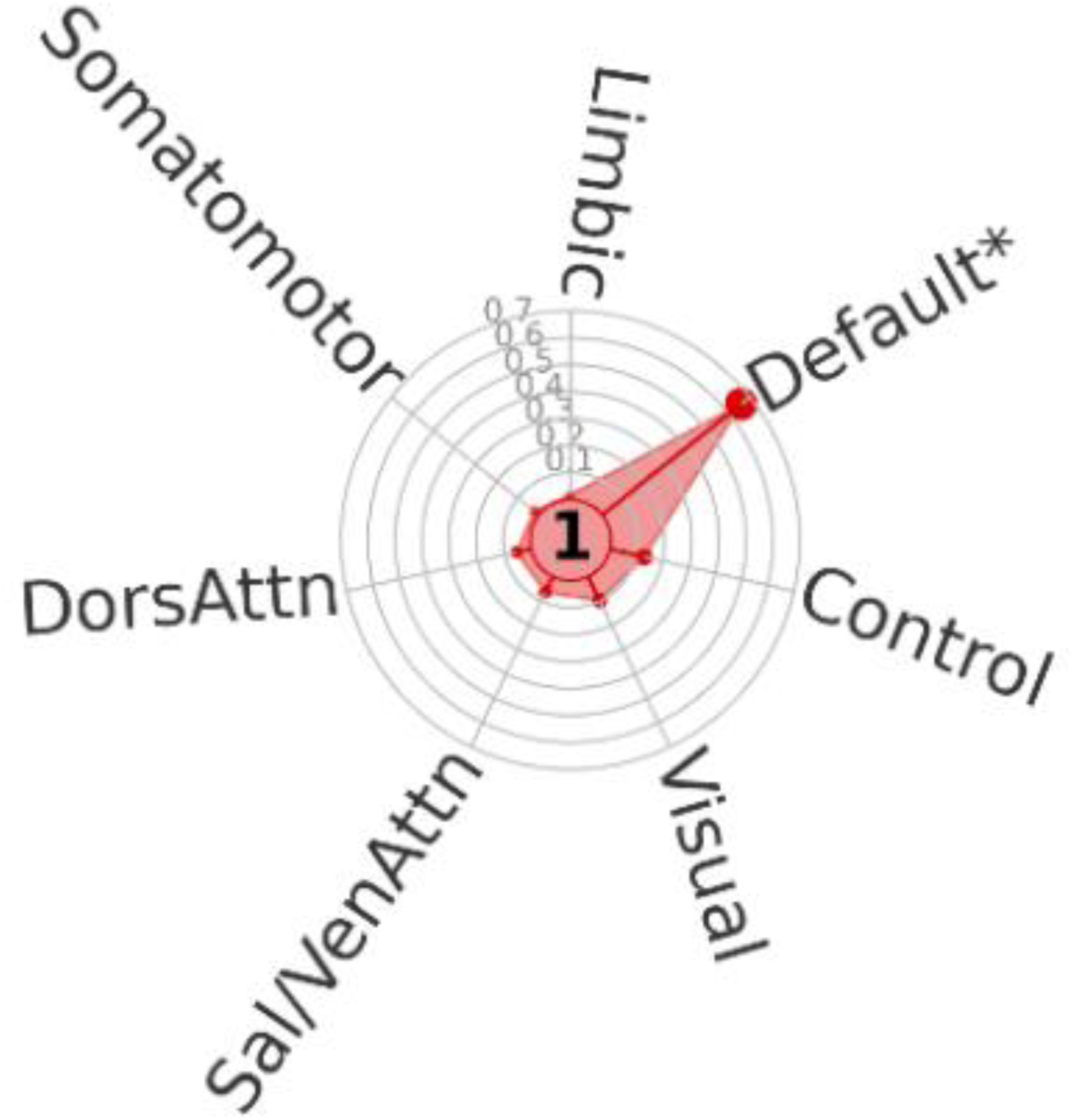
Default Mode B (DMB) overlap with “TY7” resting state networks (Kong et al., 2024).

